# Brainstem premotor circuits transform dental sensory input into coordinated jaw opening

**DOI:** 10.64898/2026.10.14.759217

**Authors:** Ling-Yu Liu, Hankyu Lee, Nathan Wagner, Kelli Hunter, Elizabeth Lee, Brian S.C. Constantinescu, Leen Dakhilalla, Amy Chang, Bo Duan, Joshua J. Emrick

## Abstract

Protective actions require the rapid transformation of damage into coordinated activity of motor pools, yet the brainstem circuits implementing this operation remain poorly defined. Here, we use the tooth-trigeminal system to identify molecularly specified pathways linking intradental sensory input to opposing jaw motor pools. Optogenetic activation of tooth-associated sensory afferents evoked rapid, stimulus-locked jaw-opening. Intradental afferents formed functional monosynaptic connections with Pdyn+ neurons in the 5Sol transition zone and Calb2+ neurons in the 5Tr transition zone, identifying these regions as sensory-recipient premotor interfaces. Predominantly excitatory Pdyn+ 5Sol neurons provided monosynaptic input to anterior digastric motor neurons, whereas predominantly inhibitory Calb2+ 5Tr neurons provided bilateral monosynaptic input to masseter motor neurons. Activation of Pdyn+ 5Sol neurons was sufficient to evoke jaw opening. Together, these findings define complementary brainstem pathways positioned to couple jaw-opening drive with inhibition of antagonistic jaw-closing output, revealing a withdrawal-like sensorimotor computation adapted to cranial biomechanics.

## Introduction

Studies of spinal sensorimotor circuits controlling limb movement (particularly proprioceptive pathways) have established fundamental principles by which defined sensory afferents engage interneurons and motor neurons to generate coordinated output. By comparison, the cellular and synaptic mechanisms that transform nociceptive input into context-dependent protective actions remains less completely resolved. Classical studies established that limb withdrawal recruits excitatory and inhibitory interneuron pathways that activate flexor muscles, suppress antagonistic extensors and engage crossed pathways for postural support (Forssberg et al., 1977). How defined sensory inputs are distributed across molecularly specified interneuron populations to generate these coordinated responses remains an important question. This gap is particularly pronounced in cranial sensorimotor systems, where brainstem circuits must coordinate opposing motor pools on both sides of mechanically coupled structures.

Anatomical and functional studies have identified the parvocellular and intermediate reticular formations as major premotor regions controlling jaw, tongue, facial and swallowing-related movements (Ugolini, 1995; Fay and Norgren, 1997; Travers and Rinaman, 2002; Stanek et al., 2014; Guo et al., 2020; Takatoh et al., 2021), and implicated genetically defined excitatory populations in jaw movement (Dempsey et al., 2021). However, these circuits have largely been characterized from the perspective of motor output. The peripheral sensory channels that recruit them, the molecular identities of sensory-recipient premotor neurons and the synaptic pathways engaging them to opposing jaw motor pools during protective behavior remain poorly defined.

The tooth-trigeminal system offers a tractable model for resolving this sensory-to-motor transformation. Teeth receive dense sensory innervation and experience substantial mechanical forces during biting and mastication and provide an anatomically restricted peripheral input whose terminals are experimentally accessed. Dental stimulation evokes a rapid and quantifiable motor response involving identifiable jaw-opening and jaw-closing muscles. Classical physiological studies established the jaw-opening reflex as a protective orofacial response in which oral or dental stimulation recruits jaw-opening muscles while suppressing jaw-closing activity (Goldberg et al., 1982; Kato et al., 1982; Nakamura et al., 1982; Lund et al., 1983). The tooth-pulp-evoked jaw-opening reflex has also served as an experimentally accessible model for investigating nociceptive processing and protective motor control across species (Mason et al., 1985). We previously showed that targeted activation of intradental sensory afferents reliably evokes rapid jaw-opening responses (Ronan et al., 2025). These observations raise a mechanistic question: how is a defined dental sensory signal routed through molecularly distinct excitatory and inhibitory premotor populations to control antagonistic jaw motor pools?

Here, we identify two complementary sensorimotor pathways linking intradental sensory input to opposing jaw motor pools. Using an afferent-centered strategy that combined genetically targeted optogenetics, viral circuit tracing, activity-dependent mapping, retrograde motor-neuron labeling, brain-slice electrophysiology and quantitative behavioral analysis, we followed intradental afferents through brainstem premotor nodes to downstream motor neurons. Intradental afferents form functional monosynaptic connections with Pdyn+ neurons in the 5Sol transition zone and Calb2+ neurons in the 5Tr transition zone. In turn, predominantly excitatory Pdyn+ 5Sol neurons provide monosynaptic input to anterior digastric motor neurons, whereas predominantly inhibitory Calb2+ 5Tr neurons provide bilateral monosynaptic input to masseter motor neurons. Optogenetic activation of Pdyn+ 5Sol neurons is sufficient to evoke rapid jaw opening. Together, these findings define a molecular and synaptic architecture through which dental sensory input recruits jaw-opening drive while providing inhibitory control over antagonistic jaw-closing output, revealing a withdrawal-like computational logic adapted for cranial motor control.

## Results

### Optogenetic activation of dental sensory afferents evokes sensitized jaw-opening responses

Previously we discovered that stimulation of the sensory innervation of the tooth elicits mandibular deflection. To further explore if this phenomenon represents a sensorimotor reflex, we created a paradigm for repetitive activation of the intradental sensory neurons while monitoring jaw opening. Here, we used low-frequency optogenetic stimulation at 0.5 Hz in *Scn10a*-Cre; LSL-ChR2 mice or control cohorts (*Scn10a*-Cre-only; LSL-ChR2-only; *Scn10a*-Cre; Ai140 LSL-EGFP mice). Intradental sensory neurons were selectively activated by positioning the optic fiber over the molar (M1; **Fig. 1a,b**). As expected, following optogenetic stimulation we observed isolated jaw-opening events in *Scn10a*-Cre; LSL-ChR2 mice (**Fig. 1d**) but not control cohorts (**Fig. 1c**).

**Fig. 1.**
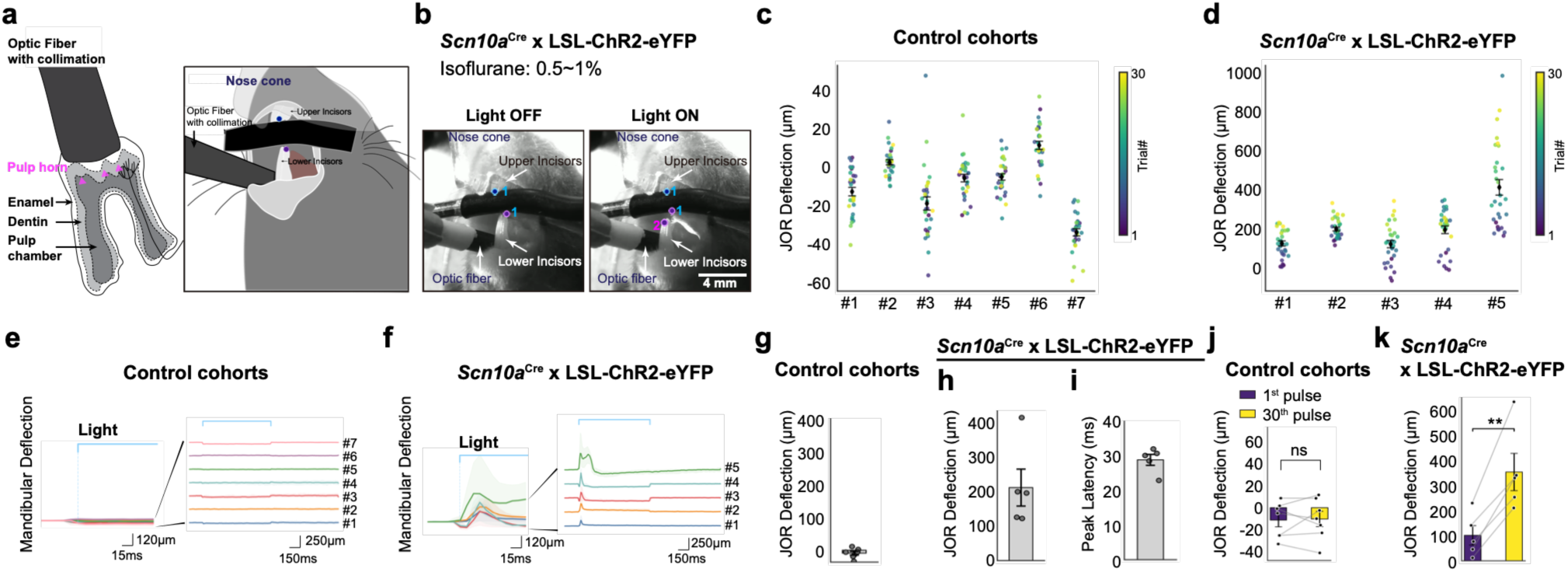
Optogenetic activation of dental sensory afferents evokes rapid and progressively enhanced jaw-opening responses. **a,** Experimental design and behavioral paradigm. *Scn10a*-Cre mice were crossed with LSL-ChR2–eYFP reporter mice to express ChR2–eYFP in *Scn10a*-expressing sensory neurons, including tooth-innervating sensory afferents. Blue-light pulses were delivered to the first maxillary molar (M1) over 1 min to activate dental sensory terminals and evoke jaw-opening responses. Schematic showing placement of the collimated optical fiber over M1. The tooth layers, from outermost to innermost, comprise enamel, dentin and the pulp chamber. Intradental sensory fibers originating in the trigeminal ganglion extend into the dentine layer. Schematic of the anesthetized preparation. Anesthesia was delivered through a nose cone, the optical fiber was positioned over M1 and the upper incisor was stabilized on a black support bar. Blue and purple landmarks indicate the upper- and lower-incisor tips, respectively; position 1 denotes the baseline light-off state. **b,** Representative images acquired before (**b,** light off) and during (**b,** light on) molar photostimulation. Position 1 marks the baseline mandibular position, and position 2 marks the displaced lower-incisor position during jaw opening. Scale bar, 4mm. **c,d,** DeepLabCut-based tracking of the upper- and lower-incisor tips and representative mandibular deflection traces over time. Stimulus-aligned mandibular deflection distances across 30 consecutive trials in stimulated *Scn10a*-Cre; LSL-ChR2–eYFP mice (**d**) and control cohorts (**c**). The dots are color-coded from the first trial (purple) to the thirtieth trial (yellow). **e,f,** DeepLabCut-based tracking of the upper- and lower-incisor tips and representative mandibular deflection traces over time. Jaw opening was quantified as the baseline-corrected change in inter-incisor distance. Videos were acquired at 107 frames per second. Representative responses were observed in stimulated *Scn10a*-Cre; LSL-ChR2–eYFP mice, n = 5 mice, (**f**); Representative responses were observed in stimulated control cohorts, n = 7 mice, (e) **g,h,** Mean jaw-opening deflection across repeated stimulation trials in stimulated *Scn10a*-Cre; LSL-ChR2– eYFP mice (**h**) and control cohorts (**g**). **i,** Mean latency from light onset to the onset of detectable mandibular deflection in stimulated *Scn10a*-Cre; LSL-ChR2–eYFP mice. **j,k,** Comparison of jaw-opening deflections evoked by the first and thirtieth light pulses in *Scn10a*-Cre; LSL-ChR2–eYFP mice (**k**) and control cohorts (**j**). Control genotypes included *Scn10a*-Cre-only, LSL-ChR2–eYFP-only and *Scn10a*-Cre; Ai140 LSL-EGFP mice. Data are presented as [mean ± SEM]. Statistical comparisons were performed using paired t-test, p-value=0.0042 (**k**), p-value=0.8476 (**j**).

We next assessed hallmarks of reflex-like properties (i.e., response reliability, latency, and adaptation) during repeated stimulation. We delivered 30 light pulses over 1 min and quantified jaw movement using the AI-based pose-estimation tool DeepLabCut (**Fig. 1e,f**). Each light pulse elicited a corresponding jaw opening event in stimulated *Scn10a*-Cre; LSL-ChR2 mice (**Fig. 1f**) with a mean jaw-opening deflection of 211.22 ± 53.37μm (mean ± SEM; 30 trials, n = 5 mice; **Fig. 1h**) compared with -9.03 ± 5.62 μm in light-stimulated control cohorts (mean ± SEM; n = 7 mice; **Fig. 1g**), and a latency of 28.94 ± 1.58 ms (mean ± SEM; n = 5 mice; **Fig. 1i**]. Negative displacement values occasionally observed in control animals reflected spontaneous jaw-clenching movements that were not time-locked to light delivery. Surprisingly, repeated stimulation resulted in a jaw-opening amplitude increase in *Scn10a*-Cre; LSL-ChR2 mice but not in the control cohorts (**Fig. 1j,k**). This progressive enhancement suggests a sensitization-like process amplifying jaw-opening output with repeated sensory activation. Importantly, optogenetic stimulation also elicited jaw-opening events in *Chrna7*-Cre; LSL-ChR2 mice, validating that activation of tooth-associated sensory afferents is sufficient to evoke this withdrawal-like motor response (**Extended Data Fig.1**, n = 4 mice). Together, the short latency, reliable stimulus locking, and stereotyped mandibular deflection establishes that jaw-opening represents a reflex response further meriting exploration of its circuitry.

### AAV-i.d. tracing reveals three brainstem targets of intradental sensory afferents

To define the sensory pathways that underlie the protective jaw-opening reflex, we first traced the central projections of intradental sensory neurons using adeno-associated viral vector-mediated intradental transduction in transgenic mouse lines (Liu et al., 2026) (**Fig. 2a, b**). To ensure rigor, we used four combinations of AAV and transgenic mouse lines to achieve labeling. Specifically, in *Scn10a*-Cre or *S100b*-Cre mice that label intradental neurons, AAV6.2-dlox-reporter was delivered into the tooth (**Fig. 2c-e**; n = 5 mice and **Extended Data Fig. 2d-f**; n = 5 mice). Conversely, in Cre-dependent reporter mice, including LSL-EGFP and/or LSL-tdTomato lines, AAV6.2-iCre was used to drive reporter expression (**Extended Data Fig. 2g-l**; n = 8 mice). Following all AAV-i.d. combinations, we detected intradental neuron somata in ipsilateral trigeminal ganglia (**Fig. 2b**).

**Fig. 2.**
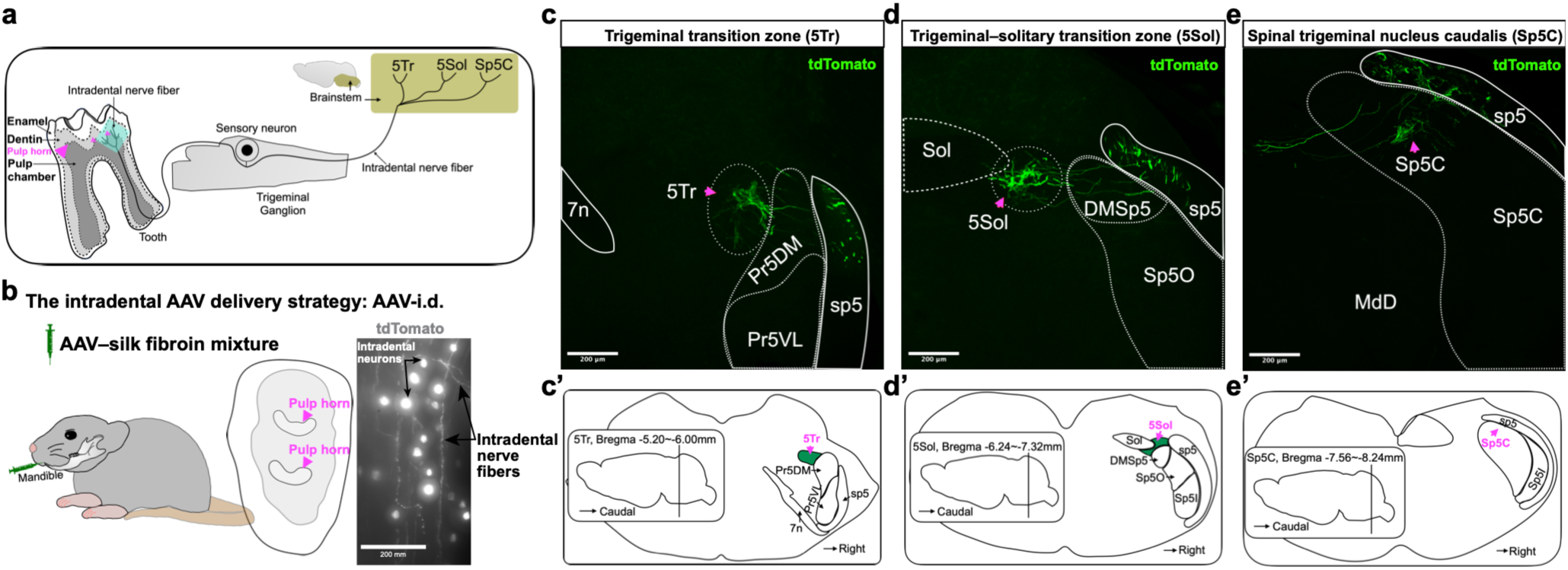
AAV-i.d. tracing identifies three brainstem targets of intradental sensory afferents. **a,** Schematic organization of the tooth–trigeminal sensory pathway. The cell bodies of tooth-innervating intradental sensory neurons reside in the trigeminal ganglion. These pseudounipolar neurons extend peripheral axons to the dental pulp and dentin and central axons to the brainstem. **b,** Intradental AAV delivery strategy (AAV-i.d.). An adeno-associated viral vector was delivered directly into the exposed pulp chamber to transduce tooth-innervating sensory neurons from their peripheral terminals. Representative image showing AAV-labeled intradental sensory fibers and the cell bodies as indicated. tdTomato-labeled fibers, shown in gray pseudocolor. Scale bar, 200 μm. (AAV-i.d. published in Liu et al. (2026), PMID: 41832951). **c-e,** Representative coronal brainstem sections showing central projections of intradental sensory afferents. AAV-dlox-tdTomato was delivered to the molars of *Scn10a-*Cre mice, resulting in tdTomato labeling of intradental sensory neurons and their central axons. tdTomato-labeled fibers, shown in green pseudocolor. Labeled afferent fibers were detected in three principal brainstem regions: the trigeminal transition zone (5Tr; **c**), trigeminal–solitary transition zone (5Sol; **d**) and spinal trigeminal nucleus caudalis (Sp5C; **e**). Arrowheads indicate representative intradental afferent projections. Scale bars, 200 μm. The labeling pattern was observed in n = 5 mice. **c′-e′,** Corresponding anatomical reference diagrams showing the locations of 5Tr (**c′**), 5Sol (**d′**) and Sp5C (**e′**) along the rostrocaudal axis. Anatomical boundaries were assigned with reference to the Allen Mouse Brain Atlas.

Based on prior work to define sensory circuits from the tooth (Greenwood and Sessle, 1976; Sessle and Greenwood, 1976; Marfurt and Turner, 1984; Hu and Sessle, 1989; Chiang et al., 2002), we focused our evaluation of the trigeminal sensory complex and surrounding brainstem structures. Our survey revealed three prominent intradental afferent projection fields based on the Allen Mouse Brain Atlas (**Fig. 2c’-e’,** n=18 mice): the trigeminal transition zone (5Tr), between the spinal trigeminal tract (sp5) and facial nerve (7N); the trigeminal-solitary transition zone (5Sol), lateral to the solitary nucleus (Sol); and the spinal trigeminal nucleus caudalis (Sp5C), with labeled projections enriched in its dorsal region (**Fig. 2c-e**). Whereas Sp5C has an established role in nociceptive processing, 5Tr and 5Sol have no ascribed functions. We observed three major intradental afferent projection domains even under sparse-labeling conditions (**Extended Data Fig. 2j-l**). Together, these results indicate that discrete brainstem targets that receive intradental sensory input with 5Sol and 5Tr representing priority candidate regions for evaluation.

### Intradental neurons send collateral projections to brainstem nuclei

Do brainstem regions associated with intradental projections receive isolated individual or common collateral inputs? We reasoned that focal stereotactic injections into individual brainstem targets would provide distinction (**Extended Data Fig. 2m**). For these experiments we relied on co-injections of Cre-dependent AAV9-dlox-GFP and cholera toxin subunit B-AF647 (CTB-647) to achieve simultaneous transduction and projection labeling with injection site visualization. As expected, we found that focal injection into the 5Sol resulted in CTB-647 labeling within the target sites (**Extended Data Fig. 2o**). Conversely, we found GFP-labeled afferent fibers associated with all projection domains (5Tr, 5Sol, and Sp5C) (**Extended Data Fig. 2n-p**; n = 7 mice). Following a second round of focal injections into the 5Tr nucleus, we again observed labeling of GFP-labeled projections to all three sites (**Extended Data Fig. 3b**; n = 3 mice). These data directly demonstrate that *Scn10a*-associated innervation of the 5Tr, 5Sol, and Sp5C are linked and suggest that intradental sensory neurons distribute input via collateral projections.

### Brainstem injections reveal molar-innervating intradental afferents

We next sought to determine the peripheral origins of associated sensory fibers (**Fig. 3b, and Extended Data Fig. 2n-p**). To this end, after stereotactic injections we evaluated the ipsilateral trigeminal ganglia and molars. As predicted by our retrograde labeling experiments, AAV-dlox-GFP into the 5Sol nucleus of *Scn10a*-Cre mice produced GFP-positive cell bodies within the ipsilateral trigeminal ganglia across the dorsal and ventral surfaces (**Fig. 3c**). Furthermore, evaluation of ipsilateral molars revealed GFP-positive fibers in the dental pulp extending into the inner dentin (**Fig. 3d-i’**). This localization was observed in both single optical sections and projections of short z-stacks (**Fig. 3d-e’**), indicating that it was not an artifact of image projection. Negative-control sections without primary antibody (**Extended Data Fig. 3e-e’**) lacked GFP-immunoreactivity. Of note, unilateral M1, M2, and M3 molars featured GFP-positive sensory innervation of pulp and dentin (n = 3 mice). Following focal injections into the 5Tr nucleus, we again observed consistent localization of GFP-positive sensory innervations in each pulp and dentin (**Extended Data Fig. 3d-d’ and f-f’**; n = 3 mice). These findings provide peripheral validation that the 5Tr or 5Sol-injections captured tooth-innervating sensory afferents targeting the tooth pulp and dentin. Together with the distributed brainstem labelling pattern, these findings support a model in which 5Sol and 5Tr receive collateral input from intradental sensory neurons.

**Fig. 3.**
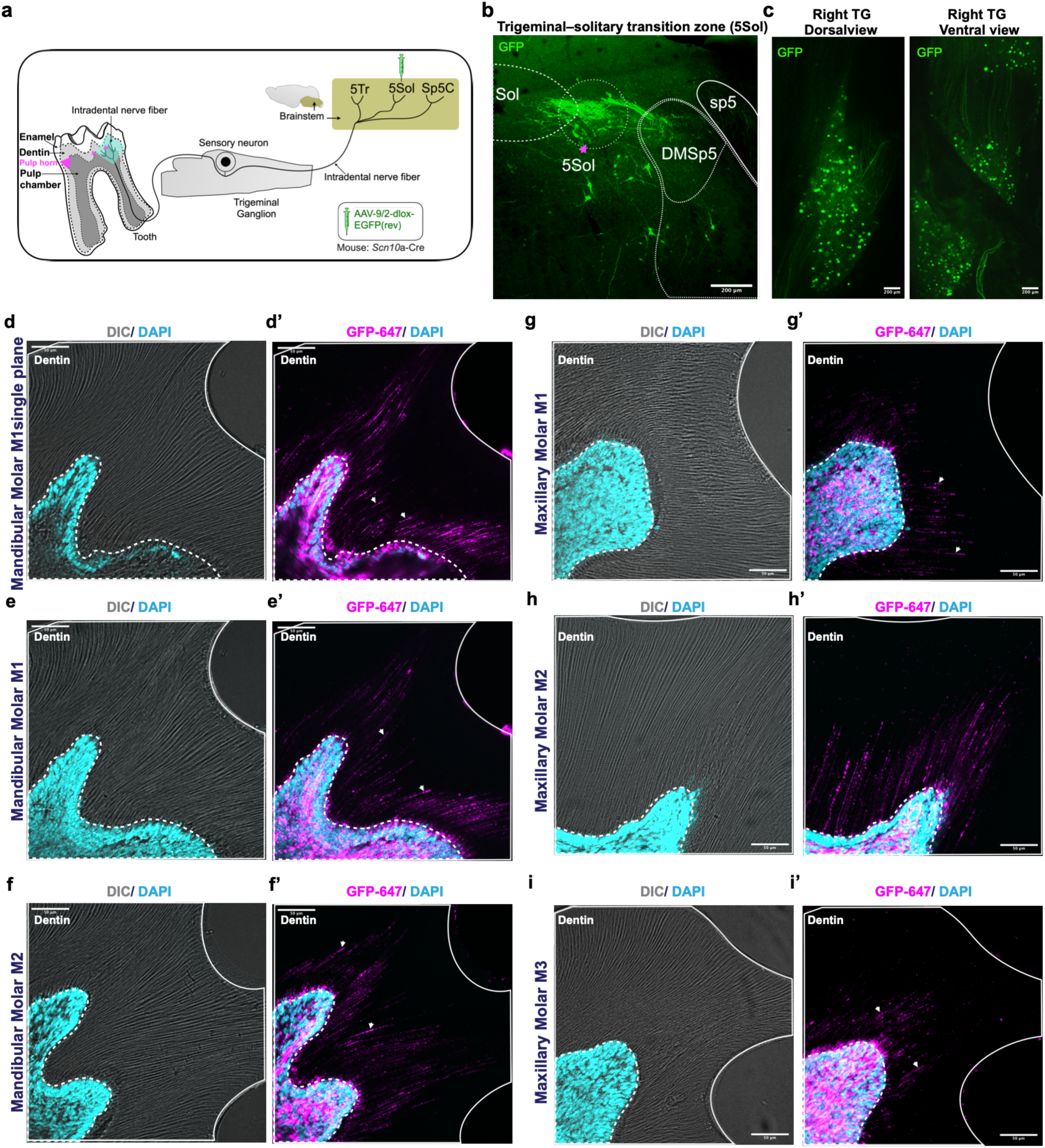
5Sol-targeted brainstem labeling identifies molar-innervating intradental sensory afferents. **a,** Schematic of the brainstem-targeted tracing strategy used to determine whether sensory afferents associated with the 5Sol projection domain innervate the molars. AAV-dlox-EGFP was stereotactically injected into the 5Sol of *Scn10a*-Cre mice, enabling Cre-dependent labeling of Scn10a-expressing neurons and their processes. **b,** Representative coronal brainstem section showing the 5Sol injection site and GFP-labeled sensory fibers following stereotactic delivery of AAV-dlox-EGFP. GFP-labeled fibers were also detected in 5Tr and Sp5C consistent with the distributed projection pattern identified by intradental tracing. **c,** Representative dorsal and ventral views of the trigeminal ganglion showing GFP-labeled sensory neuron cell bodies after 5Sol-targeted AAV delivery. GFP is shown in green. The labeling pattern was observed in n = 3 mice. **d-i′,** Representative images of decalcified molar sections showing GFP-labeled sensory fibers within the dental pulp and extending toward the dentin (white arrows). GFP signal is displayed in magenta pseudocolor, and DAPI-labeled nuclei are shown in cyan. White dashed lines delineate the pulp–dentin boundary. Labeled fibers were detected in the mandibular first molar (M1; **d-e′**) and second molar (M2; **f,f′**), as well as in the maxillary M1 (**g,g′**), M2 (**h,h′**) and third molar (M3; **i,i′**). Unilateral molar labeling was reproducibly observed in n = 3 mice. Scale bars, 200 μm (**b,c**) and 50 μm (**d–i′**).

### Postsynaptic neurons in 5Tr and 5Sol receive monosynaptic input from intradental afferents

While Sp5C relays nociception input within the trigeminal complex, 5Sol and 5Tr represent nuclei with heretofore unknown function. We reasoned that defining the molecular identity of neurons within the 5Sol and 5Tr would provide basis for functional evaluation and establish their independence from other trigeminal brainstem structures. To this end, we first evaluated *in situ* hybridization datasets from the Allen Mouse Brain Atlas (https://mouse.brain-map.org; Lein et al., 2007). Among genes known to be expressed within the brainstem (Dowsett et al., 2021; Gannot et al., 2024), we restricted our interest to those focally expressed in 5Sol/5Tr, identifying candidate molecular markers (*Calb2, Gad1, SSt, Tac1, Nts, Lhx1, Dlk1, Pdyn, Slc17a7*). We next validated candidate expression using *in situ* hybridization with afferent labeling (*Scn10a*-Cre; LSL-GFP or AAV-i.d.). These two approaches resulted in two candidate populations: Calbindin 2-positive (Calb2+) neurons and Prodynorphin-positive (Pdyn+) neurons. *Calb2* expression was enriched in the trigeminal transition zone (5Tr; **Fig. 4a**), whereas *Pdyn* expression was enriched in the trigeminal-solitary transition zone (5Sol; **Fig. 4b**). Both markers delineated neurons that were spatially-distinct from surrounding nuclei including the principal sensory and spinal trigeminal subnuclei. Together, our findings nominate Calb2+ 5Tr neurons and Pdyn+ 5Sol neurons as candidate postsynaptic partners of intradental sensory afferents.

**Fig. 4.**
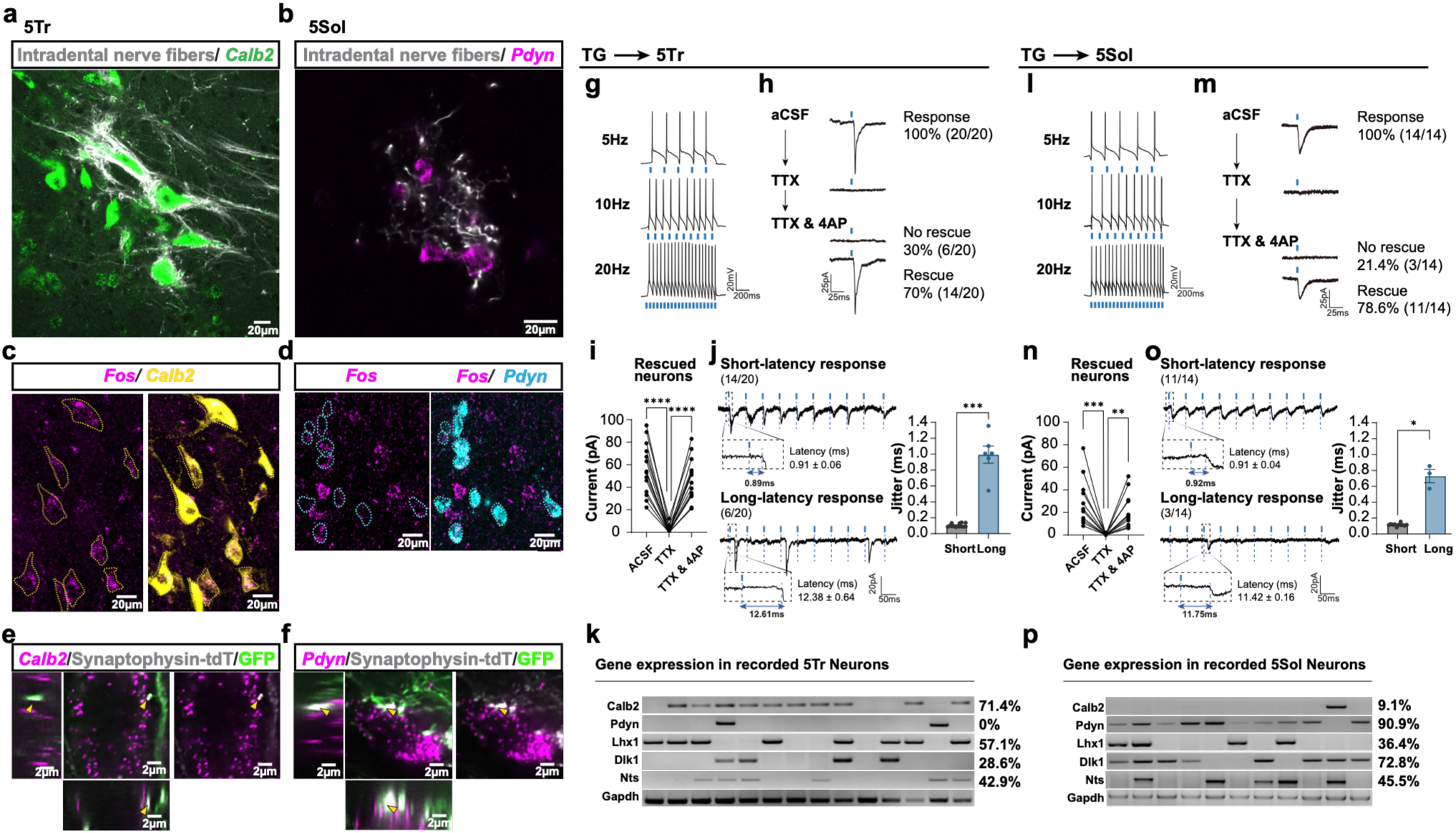
Intradental sensory afferents form monosynaptic connections with Calb2+ 5Tr and Pdyn+ 5Sol neurons. **a, b,** Identification of candidate postsynaptic populations in brainstem regions receiving intradental sensory input. Representative coronal sections show intradental afferent fibers (gray) in close proximity to Calb2+ neurons in 5Tr (green; **a**) and Pdyn+ neurons in 5Sol (magenta; **b**). Scale bars, 20 μm. Representative patterns were observed in n = 5 mice. **c, d,** Activity-dependent recruitment of candidate postsynaptic populations during optogenetically evoked jaw opening. Representative multiplex in situ hybridization images show Fos expression in Calb2+ 5Tr neurons (**c**) and Pdyn+ 5Sol neurons (**d**) after optogenetic activation of intradental sensory afferents. Fos transcripts are shown in magenta, *Calb2* transcripts in yellow and *Pdyn* transcripts in cyan. Scale bars, 20 μm. Representative patterns were observed in n = 5 mice. **e, f,** Anatomical apposition of intradental presynaptic terminals and candidate postsynaptic neurons. AAV-iCre was delivered intradentally to Ai140 LSL-EGFP; Ai34 LSL-Synaptophysin-tdTomato reporter mice, labeling intradental afferent axons with EGFP and their presynaptic specializations with Synaptophysin-tdTomato. High-resolution orthogonal views show Synaptophysin-positive intradental terminals (gray) closely apposed to Calb2+ neurons in 5Tr (magenta; **e**) and Pdyn+ neurons in 5Sol (magenta; **f**). Intradental afferent projections are shown in green. Arrowheads indicate representative sites of close apposition in the XY, XZ and YZ planes. Scale bars, 2 μm. Representative patterns were observed in n = 3 mice. **g, l,** Functional connectivity between intradental sensory afferents and postsynaptic neurons in the 5Tr and 5Sol.Representative postsynaptic responses recorded from neurons in the 5Tr (**g**) and 5Sol (**l**) during photostimulation of ReaChR-expressing intradental sensory afferent terminals. Blue bars indicate 1-ms photostimulation at ∼560 nm. **h,m,** Pharmacological assessment of light-evoked EPSCs in 5Tr (**h**) and 5Sol (**m**) neurons. All recorded neurons exhibited light-evoked responses under baseline aCSF conditions. TTX abolished the responses, and subsequent application of 4-AP in the continued presence of TTX restored light-evoked EPSCs in a subset of neurons. The numbers of 4-AP-rescued and non-rescued neurons are indicated in the figure. **i,n,** Quantification of light-evoked EPSC amplitudes in 4-AP-rescued neurons under baseline aCSF, TTX, and TTX plus 4-AP conditions in the 5Tr (**i**) and 5Sol (**n**). **j,o,** Temporal properties of light-evoked responses during repetitive photostimulation. Short-latency responses were observed exclusively in neurons whose responses were rescued by 4-AP, whereas long-latency responses were observed exclusively in non-rescued neurons. Latency jitter was significantly lower in the short-latency population than in the long-latency population. In the 5Tr, jitter was 0.103 ± 0.006 ms (n = 14) and 0.994 ± 0.108 ms (n = 6) for short- and long-latency responses, respectively (**j**). In the 5Sol, jitter was 0.116 ± 0.005 ms (n = 11) and 0.730 ± 0.084 ms (n = 3), respectively (**o**). **k,p** Post hoc molecular identification of recorded neurons by single-cell RT-PCR. In the 5Tr, 71.4% of directly connected neurons expressed *Calb2* (10/14), whereas 90.9% of directly connected 5Sol neurons expressed *Pdyn* (10/11). Recorded cells were screened for *Calb2, Pdyn, Lhx1, Dlk1,* and *Nts*, with *Gapdh* serving as an internal positive control. Recordings were obtained from 5 mice for TG→5Tr experiments and 4 mice for TG→ 5Sol experiments. Scale bars are indicated in the traces.

We next asked whether candidate populations are recruited during tooth-evoked jaw-opening responses. Using optogenetics (**Fig. 1**), we activated intradental sensory afferents and assessed activity-dependent *Fos* expression. Following stimulation via the molar, we used multiplex *in situ* hybridization to detect *Fos*, *Calb2*, and *Pdyn* transcripts and found that stimulation of the molar revealed *Fos* expression in the ipsilateral brainstem (**Fig. 4c-c’, d-d’, Extended Data Fig. 4**) in Calb2+ neurons in 5Tr (**Fig. 4c-c’**) and Pdyn+ neurons in 5Sol (**Fig. 4d-d’**). These findings indicate the ipsilateral recruitment of both molecularly-defined populations during optogenetically evoked jaw-opening behavior and identity them as candidate postsynaptic targets of intradental sensory afferents.

We next asked whether Calb2+ and Pdyn+ neurons are anatomically poised to receive intradental sensory input. Specifically, we sought to determine if presynaptic machinery of intradental afferents approximates putative postsynaptic populations. For these experiments we again relied on AAV-i.d. to perform dual fluorescent labeling (AAV-Cre into Ai140 LSL-GFP; Ai34 LSL-Synaptophysin-tdTomato mice) to express a tdTomato tagged presynaptic marker, Synaptophysin, as well as generalized cytosolic GFP expression for anatomical context. Following AAV-i.d., synaptophysin-positive intradental afferent terminals were adjacent to Calb2+/Tr neurons as visualized in orthogonal XY, YZ, and XZ planes (**Fig. 4e**). In parallel, synaptophysin-positive intradental afferent terminals were near Pdyn+/5Sol neurons in orthogonal views (**Fig. 4f**). Together, our results indicate that Calb2+/5Tr neurons and Pdyn+/5Sol neurons are candidate postsynaptic targets of intradental sensory afferents.

Finally, we directly evaluated functional connectivity between intradental sensory afferents and neurons in the 5Tr and 5Sol using optogenetic-guided brain slice electrophysiology. Intradental sensory neurons were transduced by AAV-i.d. to drive ChR2 expression in their central afferent projections followed by whole-cell recordings from either 5Tr or 5Sol during optogenetic stimulation. Here, photostimulation reliably evoked postsynaptic responses in recorded neurons in 5Sol or 5Tr indicating connectivity (**Fig. 4g,l**). To determine whether responses reflected direct monosynaptic input from intradental sensory afferents, we examined light-evoked EPSCs under aCSF, TTX, and TTX+4-AP conditions. All recorded neurons exhibited light-evoked EPSCs under baseline aCSF conditions. While TTX abolished these responses, subsequent application of 4-AP with TTX restored light-evoked EPSCs in most neurons in both the 5Tr and 5Sol (**Fig. 4h-i,m-n**), suggesting direct monosynaptic transmission from intradental sensory afferents to Pdyn+/5Sol or Calb+/5Tr neurons.

We next examined the temporal properties of light-evoked responses during repetitive photostimulation. Two clearly distinct response patterns emerged. Neurons whose responses were rescued by 4-AP consistently exhibited short-latency responses with very low trial-to-trial jitter, whereas neurons that were not rescued by 4-AP exhibited long-latency responses with substantially greater jitter (**Fig. 4j,o**). In the 5Tr, mean jitter was 0.103 ± 0.006 ms (n = 14) for the short-latency population and 0.994 ± 0.108 ms (n = 6) for the long-latency population. In the 5Sol, mean jitter was 0.116 ± 0.005 ms (n = 11) and 0.730 ± 0.084 ms (n = 3) for the short- and long-latency populations, respectively. Notably, all short-latency responses were observed in neurons that exhibited TTX/4-AP rescue, whereas all long-latency responses were observed in neurons that failed to show 4-AP rescue. Thus, the short-latency, low-jitter response phenotype corresponded precisely with the pharmacological evidence further indicating direct monosynaptic connectivity.

Following electrophysiological recordings, individual neurons were collected for single-cell RT-PCR analysis of *Calb2, Pdyn, Lhx1, Dlk1,* and *Nts*, with *Gapdh* included as an internal positive control (**Fig. 4k, p**). Among the directly connected 5Tr neurons, 10/14 cells (71.4%) expressed *Calb2*, whereas none expressed *Pdyn*. In contrast, 10/11 directly connected 5Sol neurons (90.9%) expressed *Pdyn*, whereas *Calb2* was detected in only one cell (9.1%). Together, these electrophysiological and molecular analyses demonstrate that intradental sensory afferents primarily provide direct monosynaptic input to molecularly-distinct neuronal populations in the 5Tr (*Calb2*) and 5Sol (*Pdyn*).

### Pdyn+/5Sol and Calb2+/5Tr are premotor neurons that are positioned to coordinate digastric activation and masseter inhibition

Simple spinal reflexes involve three neurons in series to achieve motor outputs within tens of milliseconds (Schieppati, 1987). Given the diminutive latency of jaw opening with optogenetics, we sought to determine whether Pdyn+ and Calb2+ neurons represent premotor neurons. Considering that jaw movement is a coordinated effort by antagonistic muscles that insert on the mandible, we hypothesized that Pdyn+ and Calb2+ populations should differentially engage motor neuron pools controlling jaw-opening and jaw-closing muscles to achieve mandibular deflection during tooth-evoked jaw-opening reflex.

We first determined the general neurotransmitter identity of Pdyn+ and Calb2+ neurons using multiplex *in situ* hybridization for *Pdyn* or *Calb2* transcripts together with excitatory and inhibitory neuronal markers, *Vglut2* and *Vgat*, respectively. We reasoned that this would reveal the nature of the signal relayed by each population (i.e., excitatory or inhibitory). In the 5Sol nucleus, the majority of Pdyn+ neurons co-expressed *Vglut2* (80.5%, n = 123 neurons, n = 3 mice, **Fig. 5c**), whereas only a small fraction co-expressed *Vgat* (4.9%). In contrast, in the 5Tr nucleus, Calb2+ neurons were predominantly inhibitory as evidenced by their co-expression of *Vgat* (73.0%, n = 125 neurons, n = 3 mice, **Fig. 5j**), whereas 16.6% co-expressed *Vglut2* among analyzed neurons. These results, consistent with the molecular analyses following brainstem whole-cell recordings (**Fig. 4**), implicate Pdyn+ 5Sol neurons as primarily excitatory and Calb2+ 5Tr neurons as primarily inhibitory, hinting at distinct roles in coordinating jaw-opening motor output.

**Fig. 5.**
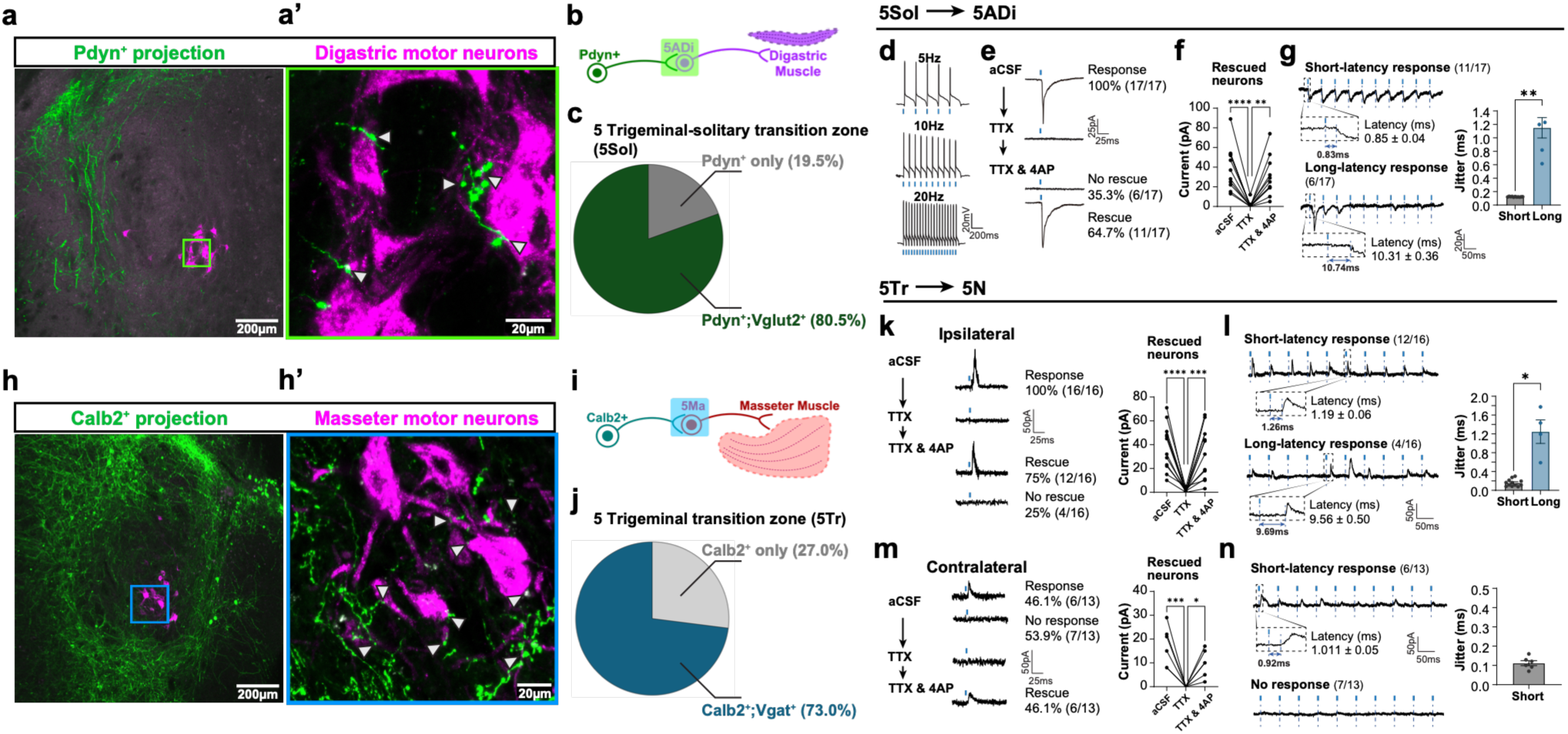
Excitatory Pdyn+ 5Sol and inhibitory Calb2+ 5Tr premotor neurons provide monosynaptic input to opposing jaw motor pools. **a,** Anatomical connectivity between Pdyn+ 5Sol neurons and anterior digastric motor neurons. Representative coronal section showing axonal projections from Pdyn+ 5Sol neurons (green) and anterior digastric motor neurons retrogradely labeled by injection of CTB-555 into the anterior digastric muscle (magenta; **a**). High-magnification view of the boxed region in **a** shows Pdyn+ axon terminals closely apposed to CTB-labeled anterior digastric motor neurons (**a′**). Scale bars, 100 μm (**a**) and 20 μm (**a′**). Representative patterns were observed in n = 5 mice. **b,** Schematic of the tracing strategy. AAV-dlox-EGFP was stereotactically injected into 5Sol of Pdyn-Cre mice to label Pdyn+ neurons and their axonal projections, while CTB-555 was injected into the anterior digastric muscle to retrogradely label its motor neurons in the anterior digastric motor pool (5ADi). **c,** Neurotransmitter identity of Pdyn+ 5Sol neurons assessed by multiplex in situ hybridization for *Pdyn* and *Vglut2* transcripts. 80.5% of Pdyn+ neurons co-expressed *Vglut2* ([99 of 123 cells]; n = 3 mice), indicating that most Pdyn+ 5Sol neurons are excitatory. **d-g,** Functional connectivity between Pdyn+ 5Sol neurons and anterior digastric motor neurons. **d,** Representative action potentials evoked in retrogradely identified anterior digastric motor neurons by photostimulation of ReaChR-expressing Pdyn+ 5Sol axon terminals at 5, 10, and 20 Hz. **e,** Representative light-evoked EPSCs recorded from anterior digastric motor neurons under baseline aCSF, TTX, and TTX plus 4-AP conditions. All recorded neurons responded to photostimulation under baseline conditions (17/17). TTX abolished the responses, and subsequent application of 4-AP in the continued presence of TTX restored EPSCs in 11 of 17 neurons (64.7%), whereas 6 of 17 neurons (35.3%) showed no rescue. **f,** Quantification of light-evoked EPSC amplitudes in the 4-AP-rescued neurons under aCSF, TTX, and TTX plus 4-AP conditions. **g,** Temporal analysis of light-evoked responses during repetitive photostimulation. All 4-AP-rescued neurons exhibited short-latency responses (11/17; latency, 0.85 ± 0.04 ms) with low trial-to-trial jitter, whereas all non-rescued neurons exhibited long-latency responses (6/17; latency, 10.31 ± 0.36 ms) with significantly greater jitter. Recordings were obtained from 6 mice. Blue bars indicate 1-ms photostimulation at ∼560 nm. Electrophysiological scale bars are indicated in the traces. **h,** Anatomical connectivity between Calb2+ 5Tr neurons and masseter motor neurons. Representative coronal section showing axonal projections from Calb2+ 5Tr neurons (green) and masseter motor neurons retrogradely labeled by injection of CTB-555 into the masseter muscle (magenta; **h**). High-magnification view of the boxed region in **h** shows Calb2+ axon terminals closely apposed to CTB-labeled masseter motor neurons (**h′**). Scale bars, 100 μm (**h**) and 20 μm (**h′**). Representative patterns were observed in n = 5 mice. **i,** Schematic of the tracing strategy. AAV-dlox-EGFP was stereotactically injected into 5Tr of *Calb2*-Cre mice to label Calb2+ neurons and their axonal projections, while CTB-555 was injected into the masseter muscle to retrogradely label masseter motor neurons in the masseter motor pool (5Ma). **j,** Neurotransmitter identity of Calb2+ 5Tr neurons assessed by multiplex in situ hybridization for *Calb2* and *Vgat* transcripts. 73.0% of Calb2+ neurons co-expressed *Vgat* ([91 of 125 cells]; n = 3 mice), indicating that most Calb2+ 5Tr neurons are inhibitory. **k-n,** Functional connectivity between Calb2+ 5Tr neurons and ipsilateral and contralateral masseter motor neurons. Cre-dependent ReaChR was expressed unilaterally in Calb2+ 5Tr neurons, and masseter motor neurons were retrogradely identified by CTB-555 labeling. **k,** Representative light-evoked IPSCs recorded from ipsilateral masseter motor neurons under baseline aCSF, TTX, and TTX plus 4-AP conditions. All ipsilateral neurons responded under baseline conditions (16/16). TTX abolished the responses, and subsequent application of 4-AP restored IPSCs in 12 of 16 neurons (75%), whereas 4 of 16 neurons (25%) showed no rescue. Quantification of light-evoked IPSC amplitudes in the rescued neurons is shown. **l,** Temporal analysis of ipsilateral responses during repetitive photostimulation. All rescued neurons exhibited short-latency responses (12/16; latency, 1.19 ± 0.06 ms) with low jitter, whereas all non-rescued neurons exhibited long-latency responses (4/16; latency, 9.56 ± 0.50 ms) with significantly greater jitter. **m,** Representative light-evoked IPSCs recorded from contralateral masseter motor neurons. Photostimulation evoked responses in 6 of 13 neurons (46.1%), whereas 7 of 13 neurons (53.9%) showed no response. TTX abolished the light-evoked IPSCs in responsive neurons, and subsequent application of 4-AP restored the responses in all 6 responsive neurons. Quantification of light-evoked IPSC amplitudes in these neurons is shown. **n,** Temporal analysis of contralateral responses during repetitive photostimulation. All responsive, 4-AP-rescued neurons exhibited short-latency responses (6/13; latency, 1.011 ± 0.05 ms) with low trial-to-trial jitter, whereas the remaining 7 neurons showed no light-evoked response. Recordings were obtained from 6 mice. Blue bars indicate 1-ms photostimulation at ∼560 nm. Electrophysiological scale bars are indicated in the traces.

To determine the putative downstream motor neurons of Pdyn+ neurons, we performed intersectional labeling to mark premotor inputs and postsynaptic motor neurons. First, we performed brainstem-targeted labeling by stereotactically-injecting AAV-dlox-GFP into the 5Sol nucleus of *Pdyn*-Cre mice. The specificity of the *Pdyn*-Cre mouse line has been validated by demonstrating overlap between Cre expression and *Pdyn* mRNA (Krashes et al., 2014). In the same animals, we injected CTB-555 into the anterior digastric muscle to retrogradely-label anterior digastric motor neurons in the brainstem 5ADi motor pool. CTB-labeled anterior digastric motor neurons co-expressed ChAT, confirming their identity as motor neurons (Armstrong et al., 1983; Houser et al., 1983). High-resolution imaging revealed Pdyn+ axonal projections were near CTB-labeled anterior digastric motor neurons (**Fig. 5a**), suggesting that Pdyn+ neurons are anatomically positioned to communicate with these effector neurons.

We next directly tested whether Pdyn+ 5Sol neurons form functional synaptic connections with anterior digastric motor neurons using optogenetic-guided brain slice electrophysiology. Pdyn+ 5Sol neurons were transduced with AAV-dlox-ReaChR to drive Cre-dependent ReaChR expression in their axonal projections, and anterior digastric motor neurons were retrogradely identified by CTB-555 labeling. Whole-cell recordings were performed in acute brainstem slices containing the trigeminal motor region while ReaChR-expressing Pdyn+ 5Sol axon terminals were locally photostimulated. Recorded motor neurons reliably followed repetitive photostimulation at 5, 10, and 20 Hz (**Fig. 5d**).

Under baseline aCSF conditions, photostimulation evoked EPSCs in all recorded anterior digastric motor neurons (17/17). TTX abolished these light-evoked responses, and subsequent application of 4-AP in the continued presence of TTX restored EPSCs in 11/17 neurons (64.7%), whereas responses in the remaining 6 neurons (35.3%) were not rescued (**Fig. 5e, f**). These pharmacological results suggest that the rescued and non-rescued neurons receive monosynaptic and potentially polysynaptic excitatory inputs, respectively.

Consistent with this distinction, the two groups exhibited markedly different response latencies and trial-to-trial jitter. All 11 neurons exhibiting TTX+4-AP rescue showed short-latency responses (0.85 ± 0.04 ms) with low trial-to-trial jitter, whereas all 6 non-rescued neurons exhibited long-latency responses (10.31 ± 0.36 ms) with significantly greater jitter (**Fig. 5g**). Thus, the consistent association of TTX+4-AP rescue with short-latency, low-jitter responses provides convergent evidence for direct monosynaptic excitatory input from Pdyn+ 5Sol neurons to a subset of anterior digastric motor neurons.

Because Calb2+ 5Tr neurons were predominantly inhibitory, we next asked how these might facilitate jaw opening. We considered two possible circuit organizations. First, Calb2+ neurons might indirectly engage motor neuron pools that would lead to jaw opening. Alternatively, Calb2+ neurons could directly inhibit motor neuron pools that elevate the mandible (i.e., masseter motor neurons), reducing jaw-closing drive and thereby facilitating jaw opening. To distinguish these possibilities, we mapped the downstream projections of Calb2+ 5Tr neurons by stereotactically-injecting AAV-dlox-GFP into the 5Tr nucleus of *Calb2*-Cre mice with validated specificity (Allen Institute for Brain Science website (**<u>Calb2-IRES-Cre images</u>**, Taniguchi et al., 2011). As predicted, we found that the projection pattern of Calb2+ 5Tr neurons was distinct from that of Pdyn+ 5Sol neurons. Specifically, Calb2+ projections did not associate with the anterior digastric motor region, nor did they exhibit concentrated localization. Instead, Calb2+ projections were highly dispersed within the trigeminal motor subnuclei, supporting a role in direct motor neuron inhibition to suppress jaw-closing (**Fig. 5h**).

We next examined whether Calb2+ 5Tr neurons communicate jaw-closing motor neurons. To label masseter motor neurons, we injected CTB-555 into the masseter muscle while stereotactically-labeling Calb2+ 5Tr neurons with AAV-dlox-GFP in *Calb2*-Cre mice. CTB-labeled masseter motor neurons were located in the 5Ma motor pool and co-expressed ChAT, confirming their motor identity (Armstrong et al., 1983; Houser et al., 1983). High-resolution imaging revealed Calb2+ axonal projections near CTB-labeled masseter motor neurons (**Fig. 5h**), suggesting that Calb2+ 5Tr neurons may engage jaw-closing motor pathways.

We next examined functional connectivity between Calb2+ 5Tr neurons and masseter motor neurons. To this end, Calb2+ 5Tr neurons were transduced with AAV-dlox-ReaChR, and masseter motor neurons were retrogradely identified by CTB-555 labeling. We then performed optogenetically guided whole-cell recordings from ipsilateral and contralateral masseter motor neurons while photostimulating ReaChR-expressing Calb2+ 5Tr axon terminals. On the ipsilateral side, all 16 recorded masseter motor neurons exhibited light-evoked IPSCs under baseline aCSF conditions. TTX abolished these responses, and subsequent application of 4-AP in the continued presence of TTX restored IPSCs in 12/16 neurons (75%), whereas responses in the remaining 4/16 neurons (25%) were not rescued (**Fig. 5k**). Consistent with the pharmacological results, response latency and trial-to-trial jitter differed markedly between rescued and non-rescued neurons. All 12 rescued neurons exhibited short-latency responses (1.19 ± 0.06 ms) with low trial-to-trial jitter, whereas all 4 non-rescued neurons exhibited long-latency responses (9.56 ± 0.50 ms) with significantly greater jitter (**Fig. 5l**). These findings support direct monosynaptic inhibitory input from Calb2+ 5Tr neurons to most ipsilateral masseter motor neurons examined.

Given the bilateral axonal projections of Calb2+ 5Tr neurons, we next examined their functional connectivity with the contralateral masseter motor pool. In contrast to the ipsilateral population, photostimulation evoked IPSCs in 6/13 contralateral motor neurons (46.1%), whereas 7/13 neurons (53.9%) showed no detectable response under baseline aCSF conditions (**Fig. 5m**). In all six responsive neurons, TTX abolished the light-evoked IPSCs and subsequent application of 4-AP in the continued presence of TTX restored the responses. These neurons also exhibited short-latency responses (1.011 ± 0.05 ms) with low trial-to-trial jitter (**Fig. 5n**). These findings demonstrate that Calb2+ 5Tr neurons establish direct monosynaptic inhibitory connections with a subset of contralateral masseter motor neurons.

Together, these anatomical and functional experiments reveal a coordinated premotor circuit architecture through which dental sensory input can promote jaw opening ipsilaterally while simultaneously regulating opposing jaw-closing activity bilaterally.

### Pdyn+ 5Sol premotor neurons are sufficient to evoke jaw opening

Because Pdyn+ 5Sol neurons are predominantly excitatory and receive monosynaptic input from intradental sensory afferents. Logically, we next asked whether activation of this population is sufficient to drive jaw-opening behavior. *Pdyn*-Cre mice were crossed with LSL-ChR2-eYFP reporter mice to drive Cre-dependent ChR2-eYFP expression in Pdyn+ neurons. An optic fiber was implanted above 5Sol at the following stereotactic coordinates relative to bregma: AP, -6.95 mm; ML, 1.30 mm; DV, -4.30 mm. ChR2-eYFP expression and optic fiber placement were histologically verified in **Extended Data Fig. 5**. As predicted, optogenetic activation of Pdyn+ 5Sol neurons reliably elicited jaw-opening events time-locked to individual light pulses (**Fig. 6b**). Across 30 stimulation trials, the mean jaw-opening deflection was 71.41 ± 17.88 μm in stimulated *Pdyn*-Cre; LSL-ChR2-eYFP mice ([Mean ± SEM], n = 5 mice; **Fig. 6c, d**), but not controls (*Pdyn*-Cre-only mice, n = 3 mice).

**Fig. 6.**
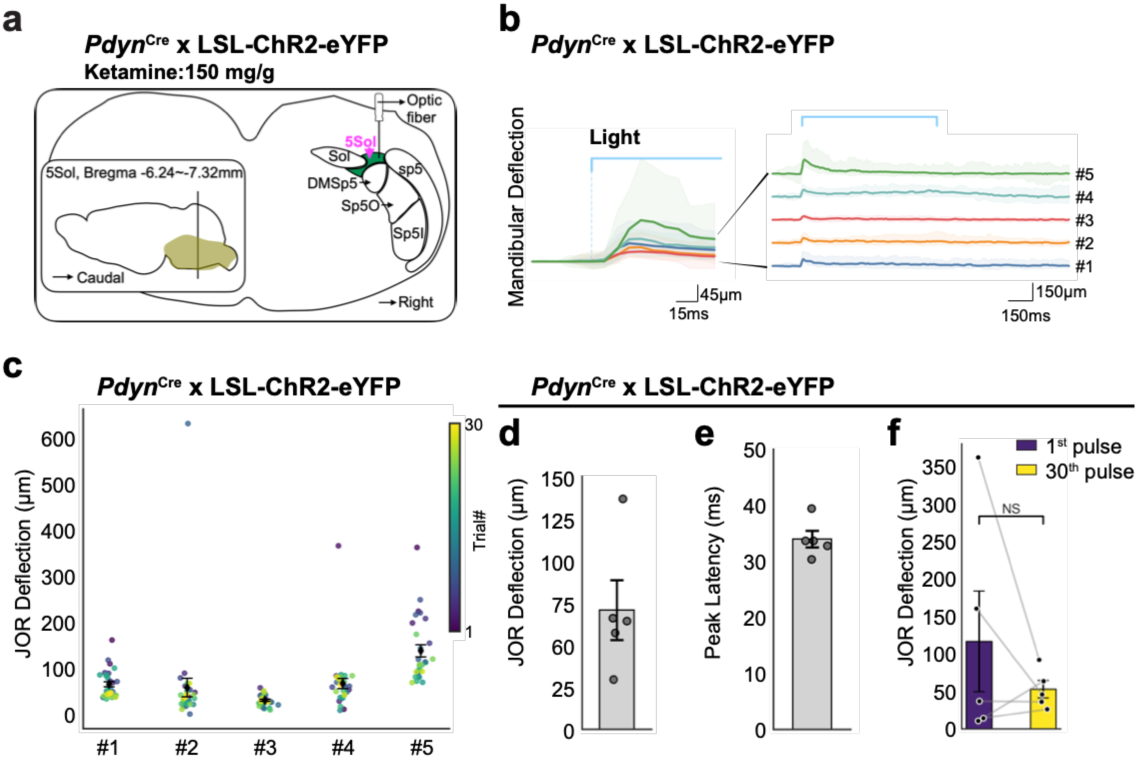
Optogenetic activation of Pdyn+ 5Sol premotor neurons is sufficient to evoke jaw opening. **a,** Experimental design. *Pdyn*-Cre mice were crossed with LSL-ChR2-eYFP reporter mice to drive Cre-dependent expression of ChR2-eYFP in Pdyn+ neurons. An optical fiber was implanted above the 5Sol at the following stereotactic coordinates relative to bregma: AP, -6.95 mm; ML, 1.3 mm; DV, -4.3 mm, and the fiber position and coordinate signs were verified in **Extended Data** Fig. 5. Blue-light pulses were delivered to the 5Sol over 1 min to activate Pdyn+ neurons and assess the resulting jaw-opening behavior. The schematic shows optical-fiber placement. **b,** DeepLabCut-based tracking of the upper- and lower-incisor tips and a representative mandibular deflection trace during photostimulation. Jaw opening was quantified as the baseline-corrected change in inter-incisor distance. Behavioral videos were acquired at 107 frames per second. **c,** Stimulus-aligned mandibular deflection traces across 30 consecutive photostimulation trials in *Pdyn*-Cre; LSL-ChR2–eYFP mice. Traces are color-coded from the first trial (purple) to the thirtieth trial (yellow). **d,** Mean jaw-opening deflection across repeated photostimulation trials in *Pdyn*-Cre; LSL-ChR2-eYFP mice. **e,** Latency from light onset to the onset of detectable mandibular deflection. Jaw opening became detectable approximately 34 ms after light onset, corresponding to three to four frames at the video acquisition rate of 107 frame/s. **f,** Comparison of jaw-opening deflections evoked by the first and thirtieth light pulses. Response amplitudes did not differ significantly across repeated activation of Pdyn+ 5Sol neurons. Data are presented as [mean ± SEM]. Statistical comparisons were performed using paired t-test, p-value=0.3359 (**f**); n = 5 mice.

In these experiments, activation of Pdyn+ 5Sol neurons was rapidly coupled to mandibular motor output [34.21 ± 1.50 ms (mean ± SEM; n = 5 mice; **Fig. 6e**)]. Notably, in direct departure from peripheral sensory-afferent stimulation, no significant change in Pdyn+ evoked response was detected between the first and the final trials (**Fig. 6f**, two-tiled paired t-test, P value =0.3359) indicating that peripheral activation is necessary for response facilitation. Together, these findings demonstrate that activation of Pdyn+ 5Sol premotor neurons is sufficient to evoke rapid jaw opening.

## Discussion

Protective behaviors require that sensory signals indicating tissue damage must be rapidly transformed into coordinated activity by motor pools. Classical physiological studies established that protective withdrawal depends on excitation of muscles that move the body away from a stimulus together with inhibition of their antagonists (Schouenborg et al., 1990; Levinsson et al., 1999). In the orofacial system, this computation must accommodate the distinctive biomechanics of the mandible, which requires coordinated control of opposing muscles acting on a single, bilaterally coupled structure (Kidokoro et al., 1968; Sherrington, 1910; Türker et al., 1994). Although tooth-evoked jaw opening and broad orofacial premotor territories have long been recognized, heretofore the molecular and synaptic pathways that route a defined dental sensory input to opposing jaw motor pools have not been defined.

Our work bridges this gap by defining a sensory-to-premotor-to-motor architecture for protective jaw opening. Intradental sensory afferents form functional monosynaptic connections with predominantly excitatory Pdyn+ neurons in 5Sol and predominantly inhibitory Calb2+ neurons in 5Tr. Pdyn+ 5Sol neurons, in turn, provide monosynaptic excitatory input to anterior digastric motor neurons, and their optogenetic activation is sufficient to move the jaw downward. Conversely, Calb2+ 5Tr neurons provide monosynaptic inhibitory input to masseter motor neurons bilaterally. By identifying molecularly defined neuronal populations and direct synaptic connections positioned to couple jaw-opening drive with inhibition of antagonistic jaw-closing output we advance our understanding beyond prior physiological descriptions of reciprocal jaw-muscle control (**Fig. 7**).

**Fig. 7.**
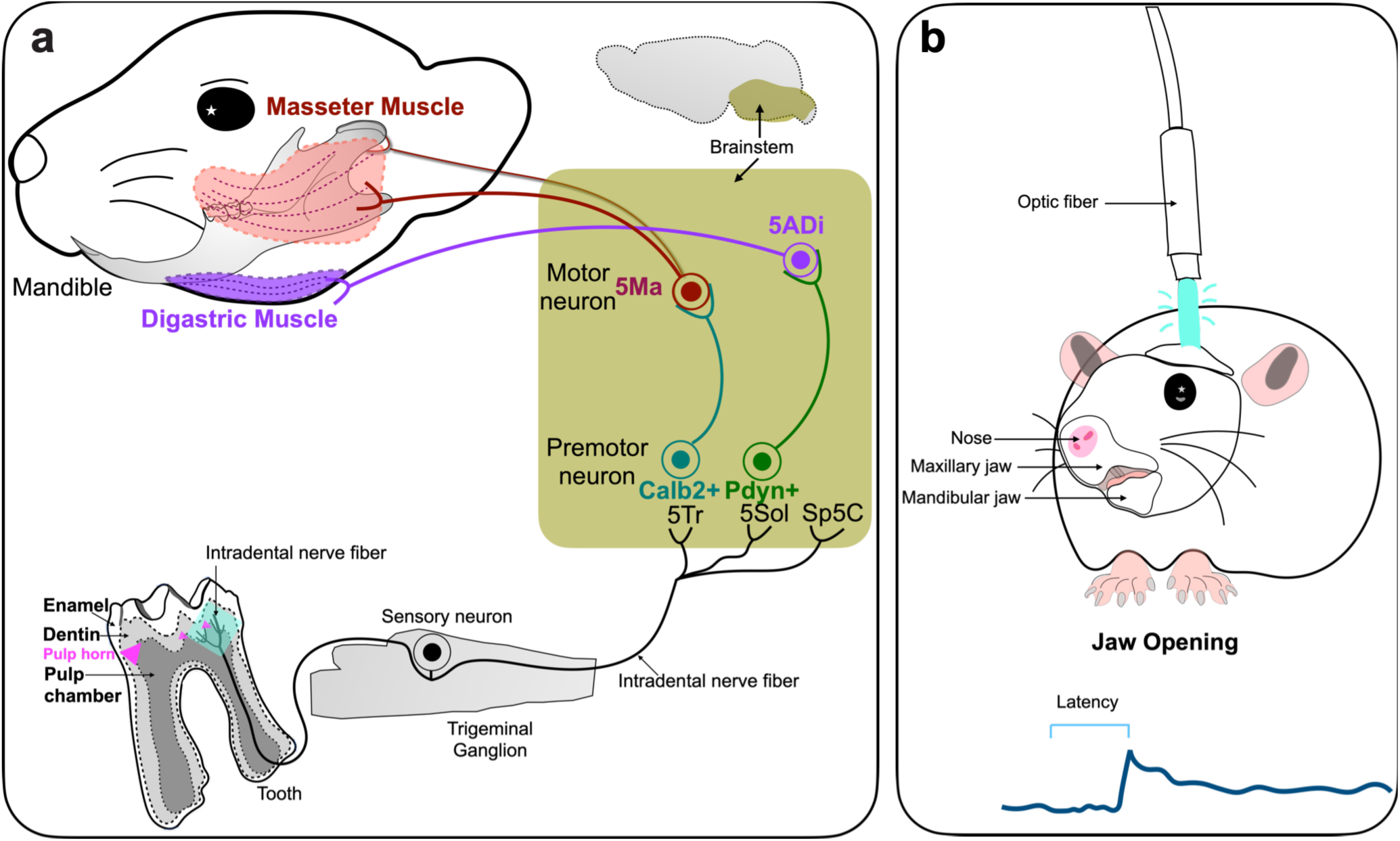
A dual premotor circuit transforms dental sensory input into coordinated jaw opening. **a,** Schematic model of the peripheral-to-brainstem circuit linking intradental sensory afferents to opposing jaw motor pathways. Intradental afferents form functional monosynaptic connections with two anatomically and molecularly distinct populations: Pdyn+ neurons in 5Sol and Calb2+ neurons in 5Tr. These populations engage complementary premotor pathways. Predominantly excitatory Pdyn+ 5Sol neurons provide monosynaptic input to anterior digastric motor neurons (5ADi), whereas predominantly inhibitory Calb2+ 5Tr neurons provide monosynaptic input to masseter motor neurons (5Ma). **b,** Optogenetic activation of Pdyn+ 5Sol neurons is sufficient to evoke jaw opening. Together, these findings support a circuit model in which dental sensory input is transformed into protective jaw-opening output by coupling excitation of jaw-opening motor neurons with inhibition of antagonistic jaw-closing motor neurons.

### 5Tr and 5Sol as an interface within the orofacial premotor network

Our findings refine current models of orofacial premotor organization by following intradental afferents downstream. Our afferent-centered strategy identified 5Tr and 5Sol as novel sensory-recipient transition zones with direct premotor access to jaw motor pools. Previous studies tracing backward from cranial motor pools identified the parvocellular reticular formation and intermediate reticular formations (PCRt and IRt) as other major sources of premotor inputs to neurons controlling the jaw, tongue, face, and pharynx (Ter Horst et al., 1991; Stanek et al., 2014; Dempsey et al., 2021; Takatoh et al., 2021; Wang & Takatoh 2026). Further anatomical, functional, and molecular analysis will be needed to define their relationship to neighboring PCRt and IRt territories and determine if 5Tr and 5Sol neurons access the broader orofacial premotor network, or represent isolated premotor centers.

### An ipsilaterally weighted pathway with cross-midline access

The architecture of the jaw-opening reflex implements a principle shared with spinal withdrawal pathways: excitation of a movement-producing pathway together with inhibition of its antagonist (Schouenborg et al., 1995; Tripodi M. et al., 2011; Arber S., 2017). Considering spinal and cranial circuits adapt reciprocal motor control to different biomechanical constraints, the similarity is solely computational. Here, unilateral mapping from the tooth revealed distinct lateral organizations of corresponding premotor outputs. Pdyn+ 5Sol neurons innervated the ipsilateral anterior digastric motor pool, whereas Calb2+ 5Tr neurons directly accessed masseter motor neurons bilaterally. Monosynaptic Calb2+ responses were detected in 12 of 16 ipsilateral and 6 of 13 contralateral masseter motor neurons. The bilateral organization of Calb2+ output likely reflects the mechanically coupled movement of the mandible (Mushimoto and Mitani, 1982). Direct ipsilateral Calb2+ connectivity could rapidly regulate masseter motor neurons on the side receiving dental input, whereas cross-midline projections provide a route through which unilateral sensory processing can influence the opposite motor pool (Jacquin et al., 1990; Kamogawa et al., 1994; Morquette et al., 2012). This bilateral access could contribute to the timing, gain or symmetry of jaw-closing activity, but the present experiments only establish synaptic connectivity rather than its degree of inhibition or contribution to behavioral function. Simultaneous bilateral electromyography combined with unilateral manipulation of Calb2+ 5Tr neurons will be required to determine if cumulative motor inhibition correlates with relative monosynaptic connections as well as how this cross-midline pathway contributes to protective jaw opening.

### Potential interaction with masticatory pattern-generating networks

The pathways identified here hint at how protective dental input modifies ongoing mastication. We suggest that brainstem pattern-generating networks produce rhythmic jaw opening and closing, while oral sensory signal continuously adjusts this motor program (Lund, 1991; Moore et al., 2014). Jaw closure may result in unexpected damaging forces onto the teeth that requires a rapid transition toward opening to prevent fracture. Direct activation of Pdyn+ 5Sol neurons evoked jaw opening in the absence of patterned peripheral stimulation, demonstrating that this population can recruit jaw-opening output downstream of the initial dental signal. Together with direct bilateral Calb2+ inhibition of masseter motor neurons, this architecture biases the direction of the masticatory motor system from closure toward opening. Experiments to manipulate the circuit during natural chewing will reveal whether these pathways also interrupt, reset or phase-dependently modify chewing rhythms.

### Limitations and future directions

Although Pdyn+ 5Sol activation was sufficient to evoke jaw opening, loss-of-function experiments are required to determine whether this population is necessary for native tooth-evoked responses. Similarly, the proposed role of Calb2+ 5Tr neurons in antagonist suppression is supported by their inhibitory identity and direct connectivity with masseter motor neurons but has not been established causally during behavior. The functional role of contralateral Calb2+ signaling also remains to be determined.

In summary, using the tooth-trigeminal pathway we determine how brainstem circuits rapidly reconfigure information into motor output to protect vulnerable tissues. We reveal for the first time how protective circuits are adapted to the biomechanics of a bilaterally coupled mandible. In this framework, we find that a spatially localized peripheral sensory signal diverges into complementary premotor commands that engage opposing and bilateral motor pools. Furthermore, by providing molecular and anatomical definition to this circuit, we add new depth to our understanding of both reciprocal agonist-antagonist control as well as a basis for a fundamental cranial reflex.

## Acknowledgements

We thank members of LeeT., Emrick, Duan, Kramer, LeeP. labs for your support, especially C.P.Y., B.T., H.C., R.A., P.K., K.H.U-K., X.L. for discussion and advice, and H.L., B.D., B.S. for constructive feedback on this paper. We thank Dr. Eric Rentchler and the Michigan Medicine BRCF Microscopy Core at the University of Michigan for training and coordinating access to the Nikon CSU-W1 SoRa.

## Author Contributions

Conceptualization: L.Y.L., B.D. and J.J.E.; Methodology: L.Y.L., H.L., N.W., K.H., E.L., B.S.C.C., L.D., A.C.; Software, validation, formal analysis: L.Y.L., H.L. and B.S.C.C.; Investigation, data curation, visualization, project administration and writing-original draft: L.Y.L. and H.L.; Resources: L.Y.L., B.D. and J.J.E.; Writing— review and editing: L.Y.L., H.L., N.W., K.H., B.D. and J.J.E.; Supervision: L.Y.L., B.D. and J.J.E.; Funding acquisition: L.Y.L., B.D. and J.J.E.

## Methods

### Animals

All animal procedures were approved by the University of Michigan Institutional Animal Care and Use Committee (IACUC; protocol number: PRO00012299) and were performed in accordance with relevant institutional guidelines and the National Institute of Health*-Guide for the care and use of laboratory animals*.

Both male and female mice were included in this study; the sex distribution for each experiment is reported in the corresponding source data. Mice were group housed except when single housing was required after surgery under controlled environmental conditions at 25°C, ∼50% humidity on a 12 h light/12 h dark cycle, with food and water available ad libitum access. Experiments were conducted during the light phase.

The following mouse lines were used: *Calb2-*IRES-Cre (#010774, JAX); *Pdyn*-IRES-Cre (#027958, JAX); *Scn10a*-Cre (*Nav1.8*-Cre, #036564, JAX); *S100b*-Cre (*S100b*-Cre provided by Dr. Nicholas Ryba and previously validated for labeling intradental sensory neurons in Ronan et al., 2025); Ai14 LSL-tdTomato (#007914, JAX); Ai140 LSL-EGFP (#034100, JAX); Ai34 LSL-Synaptophysin-tdTomato (#012570, JAX); Ai32 LSL-ChR2(H134R)-EYFP (#024109, JAX); and *Chrna7*-Cre (MGI # 5662228, provided by Dr. M. Rothermel and tooth sensory neuron innervation was validated in Rogers & Gahring, 2012). Genotypes were determined by the service of Transnetyx Inc. Cre-negative and/or reporter only littermates were used as controls where indicated.

For optogenetic activation of tooth-associated sensory afferents, *Scn10a*-Cre or *Chrna7*-Cre mice were crossed with LSL-ChR2(H134R)-EYFP mice. For optogenetic activation of Pdyn+ 5Sol neurons, *Pdyn*-Cre mice were crossed with LSL-ChR2(H134R)-EYFP mice. To label intradental afferent axons and presynaptic specializations (Synaptophysin-tdTomato), LSL-EGFP mice were crossed with LSL-Synaptophysin-tdTomato mice, and Cre recombination was induced by AAV-i.d of AAV-iCre, as described below. Experimental and genotypes used for each assay are reported in the corresponding methods subsection and figure legend.

### Surgical procedures

Animals were randomly assigned to experimental groups. All surgeries were performed under aseptic conditions. Injection and optical-fiber locations were verified histologically after each experiment. Animals were excluded according to prespecified criteria if viral expression or fiber placement did not encompass the intended target region.

#### Tooth/ Intradental AAV delivery (AAVi.d.)

AAV-i.d. was performed as described previously (Liu et al., 2026; PMID: 41832951). Mice were anesthetized with isoflurane delivered using a SomnoSuite low-flow anesthesia system (4.5% at 475 mL/min for initial induction; 2% at 175 mL/min for maintenance). Anesthetic depth was monitored by respiratory rate and absence of the pedal-withdrawal reflex. Animal were positioned prone, and the oral cavity was exposed using jaw and lateral retractors, and the suspension wire on the Assembled stereotax (MB1015/M, Thorlabs; HW-KIT2/M M6X1.0 Hardware Kit, Thorlabs; 344134, Acco).

Approximately 200-300 μm of the coronal cusps of the target molar was removed to expose two to three pulp horns using a ¼ round bur (HSB-401152, House Brand Dentistry) and a dental drill (C20 (BK-A08), Delanie Electric Nail Drill). A total of 2μL of an AAV-silk fibroin mixture was applied to the preparation through repeated cycles of application and drying processes. The AAV-silk mixture consisted of AAV6.2-iCre (v225-6(F129L), ETH VVF) for labeling and silk fibroin (50 mg/mL, #5154 Advanced BioMatrix) was used to retain and concentrate the viral vector at the exposed pulp surface during the application. The preparation was sealed with a flowable dental composite (N11VA, Pentron) and polymerized using a dental light (K192233, Net32/Ebay).

Depending on the experimental design, the following vectors were delivered intradentally: AAV6.2/2-dlox-tdTomato; AAV6.2/2-dlox-EGFP; AAV6.2/2-iCre. The reporter or Cre-dependent mouse lines paired with each vector are specified in the corresponding experimental section. Animals were allowed to recover for three weeks before tissue collection or subsequent experiments. Trigeminal ganglions were examined to confirm labeling of tooth-innervating sensory neurons.

#### Stereotactic viral injection into 5Tr and 5Sol

Mice were anesthetized with isoflurane using a SomnoSuite low-flow anesthesia system (3.9% at 393 mL/min for initial induction; 2.5% at 130mL/min for maintenance and secured in a stereotaxic frame (51730UD, Stoelting). Carprofen (5mg/kg, subcutaneous) (510510, MWI Animal Health) was administered before surgery and postoperatively for 10 days. Body temperature was maintained with a heating pad and during recovery from anesthesia. Anesthetic depth was monitored by respiratory rate and absence of the pedal-withdrawal reflex.

Hair on the scalp was removed, and the scalp was sterilized () before the initial incision was made to expose the skull. After leveling the skull using bregma and lambda, a small craniotomy was made over the target using a ¼ round drill bur. Viral vectors were delivered using a Neuros syringe (0.5 µL, 32 gauge, 65457-01, Hamilton).) coupled to a nanoliter injector (53311, Stoelting).

The viral vectors were injected into 5Tr at the coordinates relative to bregma at [AP: -5.975 mm; ML: +/- 1.55 mm; DV: -4.50 mm], and into 5Sol at the coordinates relative to bregma at [AP: -6.95 mm, ML: +/- 1.30 mm, DV: −4.50 mm].

The injection was administered at a flow rate of 40 nL/ min. After injection, the Syringe was left in place for 10 minutes and then withdrawn slowly over 5 minutes to minimize reflux. The incision was closed using suture (Y493G, Ethicon). Mice were allowed to recover on a heated surface before being returned to their home cage. Postoperative monitoring and analgesia were provided according to the approved institutional protocol.

#### Projection mapping of Calb2+ 5Tr and Pdyn+ 5Sol neurons

To label the axonal projections of Calb2+ 5Tr neurons, AAV9/2-dlox-tdTomato (v167-9, VVF) and/or AAV9/2-dlox-EGFP (v167-9, ETH VVF) was injected into 5Tr of *Calb2*-Cre mice.

To label the axonal projections of Pdyn+ 5Sol neurons, AAV9/2-dlox-tdTomato (v167-9, VVF) and/or AAV9/2-dlox-EGFP (v167-9, ETH VVF) was injected into 5Sol of *Pdyn*-Cre mice. Tissue was collected three weeks after viral injection.

#### Optogenetic circuit mapping from premotor neurons to jaw motor neurons

To assess connectivity between Calb2+ 5Tr neurons and masseter motor neurons, AAV9/2-dlox-ReaChR-mCitrine (v261-9, VVF) was unilaterally injected into 5Tr of *Calb2*-Cre mice. Three weeks later, CTB-555 (cholera toxin subunit B conjugated to Alexa Fluor 555, (1%, 10.0mg/mL, 2.5uL, C34776, ThermoFisher) was injected into the ipsilateral / bilateral masseter muscles to retrogradely label masseter motor neurons, 5Ma. Acute brainstem slices were prepared 3-5 days after CTB injection for electrophysiological recordings.

To assess connectivity between Pdyn+ 5Sol neurons and masseter motor neurons, AAV9/2-dlox-ReaChR-mCitrine (v261-9, VVF) was unilaterally injected into 5Sol of *Pdyn*-Cre mice. Three weeks later, CTB-555 (1%, 10.0mg/mL, 2.5uL, C34776, ThermoFisher) was injected into the anterior digastric muscle unilaterally to retrogradely label the corresponding motor neurons, 5ADi. Acute brainstem slices were prepared 3-5 days after CTB injection for electrophysiological recordings.

For all intramuscular injections, mice were anesthetized with isoflurane using a SomnoSuite low-flow anesthesia system (3.9% at 393 mL/min for initial induction; 2.5% at 130mL/min for maintenance) and secured in a stereotaxic frame (51730UD, Stoelting). CTB was delivered at 2.5 μL using (2.5 µL Microliter Syringe, 7632-01, Hamilton; 34 gauge needle, 207434, Hamilton).

#### Implantation of optic fiber

For optogenetic stimulation of 5Sol neurons, an optic fiber (200 μm core diameter; NA = 0.39, RWD-R-FOC-BL200C-39NA, RWD Life Science) was implanted above the target region at the coordinates specified for each experiment. The fiber tip was positioned 0.2 mm above the viral injection site, unless otherwise indicated. The implant was secured to the skull using two skull screws, adhesive cement and dental acrylic (339291, 339104, GC America).

Animals were allowed to recover for 10 days before behavioral testing. Optic fiber placement and viral expression were verified histologically after completion of the experiments. Animals were excluded if the fiber tip was outside the prespecified anatomical boundary, viral expression did not encompass the target region or expression extended substantially into the motor region. Behavioral experiments were performed 3weeks after injection.

### Viral constructs

The production, purification, and physical titration of the viral vectors were performed by the Viral Vector Facility (VVF) of the Neuroscience Center Zurich (ZNZ). Vectors were generated using plasmids from the VVF repository under standard protocols. All of the viral vectors listed below were purchased from the VVF.

AAV6.2/2-dlox-tdTomato (6.5 x 10E12 vg/ml, v167-6(F129L), ETH VVF)

AAV6.2/2-dlox-EGFP (5.7 x 10E12 vg/ml), v158-6(F129L), ETH VVF

AAV6.2/2-hEF1a-iCre (7.2 x 10E12 vg/ml, v225-6(F129L), ETH VVF)

AAV9/2-dlox-tdTomato (5.5 x 10E12 vg/ml, v167-9, VVF)

AAV9/2-dlox-EGFP (6.0 x 10E12 vg/ml, v158-9, ETH VVF)

AAV9/2-dlox-ReaChR_mCitrine (5.9 x 10E12 vg/m, v261-9, VVF)

### Optogenetic activation of intradental behavioral paradigm

For optogenetic activation of tooth-innervating sensory afferents, *Scn10a*-Cre mice were crossed with LSL-ChR2(H134R)-EYFP reporter mice to express ChR2(H134R)-EYFP in *Scn10a*-expressing sensory neurons. Control genotypes included *Scn10a*-Cre only, LSL-ChR2(H134R)-EYFP only and *Scn10a*-Cre; Ai140 LSL-EGFP mice. Male and female mice (8-12 weeks) were used

Mice were anesthetized with isoflurane using a SomnoSuite low-flow anesthesia system (3.9% at 393 mL/min for initial induction; 0.5-1% at 130mL/min for maintenance through a secured nose cone. Anesthetic depth was adjusted to maintain stable respiration and the absence of spontaneous jaw movements and pedal withdrawal reflex while preserving the evoked response. The upper incisor was stabilized on a black support bar to reduce head movement and provide consistent visualization of mandibular displacement. A collimated optic fiber (500 μm core diameter, NA= 0.63, Goldstone Scientific) emitting blue LED light was positioned over the right maxillary molar M1 with direct contacting the tooth. The optic fiber was coupled to a 470 ± 6 nm light source through a patch cable (Prizmatix High NA Plastic Optical Fibers, Goldstone Scientific). Light was delivered at 43.7 mW measured at the fiber tip using a device (Ophir PD300RM, MKS Instruments), corresponding to an estimated irradiance of average intensity 222.56 mW/mm^2^. Each animal received 30 stimulation trials over 1 minute at 0.5Hz and a 50% duty cycle. Optical power, fiber-to-tooth distance and illumination geometry were held constant across animals.

Optical stimulation was controlled by a TTL pulse generator (OTPG8, Doric Lenses, OTPG_8, Doric) and synchronized with behavioral video acquisition. The oral cavity was imaged using a Blackfly S camera (BFS-U3-13Y3M-C, #36-451, FLIR, Edmund Optics) at 107 fps corresponding to a temporal resolution of 9.35 ms per frame. The movement onset was interpreted at the frame-level resolution of the behavioral video.

Jaw movement captured was quantified using DeepLabCut (DeepLabCut GUI 2.3.5)-based tracking of the upper and lower incisor tip to measure the amplitude of the mandibular deflection. The Euclidean distance between incisors was calculated and aligned to TTL-defined light onset; jaw-movement latency was resolved at the 9.35 ms frame interval.

### Implantation of optic fiber in 5Sol

Male and females *Pdyn-*Cre; LSL-ChR2(H134R)-EYFP mice aged 12-15 weeks were used. Stereotactic coordinates were optimized in pilot animals and subsequently applied consistently across experimental cohorts. Mice were anesthetized with isoflurane delivered using a SomnoSuite low-flow anesthesia system (3.9% at 498 mL /min for induction; 2.5% at 130 mL /min for maintenance) and secured in a stereotaxic frame. Carprofen (5 mg/Kg, subcutaneous) was administered for analgesia. Anesthetic depth was monitored by respiratory rate and absence of the pedal-withdrawal reflex. Body temperature was maintained.

Hair on the scalp was removed, and the scalp was sterilized () before the initial incision was made to expose the skull. After leveling the skull using bregma and lambda, a small craniotomy was made over the target using a dental drill. An optic fiber (200 μm core diameter; NA = 0.39, RWD-R-FOC-BL200C-39NA, RWD Life Science) was implanted above the right 5Sol region at the coordinates relative to bregma: AP: -6.95 mm; ML: +/-1.30 mm; DV: −4.30 mm in *Pdyn*-Cre; LSL-ChR2(H134R)-EYFP mice. The fiber tip was positioned 0.2 mm dorsal to the intended neuronal target. The implant was secured to the skull using a thin layer of cyanoacrylate adhesive followed by dental cement (339291, 339104, GC America) to form a stable headcap. Animals recovered on a heated surface and were monitored until ambulatory before being returned to their home cages. Postoperative analgesia was administered according to the approved institutional protocol. Behavioral testing was performed after a recovery period of 10 days.

Optic fiber placement was verified histologically after completion of the experiments. Animals were excluded according to prespecified criteria if the fiber tip was outside the 5Sol target region, ChR2(H134R)-EYFP expression did not encompass the intended site or the implant caused substantial tissue damage.

### Optogenetic activation of Pdyn+ 5Sol neurons

For behavioral photostimulation, mice were initially anesthetized with isoflurane (3.9% at 393 mL /min) for approximately 2 min and were subsequently administered ketamine (150 mg /kg, ip) for maintenance. Isoflurane was then discontinued, and the animals were positioned in a stereotaxic frame. Anesthetic depth was monitored throughout the experiment using respiratory rate and the pedal-withdrawal reflex. Body temperature was maintained at approximately 37 °C using a heating pad.

The implanted optic fiber was coupled to a 473-nm light source through a patch cable (M95L01, ThorLabs). Light was delivered at 5.7 mW measured at the fiber tip, corresponding to an estimated irradiance of average intensity 181.44 mW/mm^2^. Each mouse received 30 stimulation trials over 1 min at 0.5 Hz with a 50% duty cycle. At 0.5 Hz, this duty cycle corresponds to 1 s of illumination followed by 1 s without illumination.

Photostimulation was controlled by a TTL pulse generator (OTPG8, Doric Lenses) and synchronized with behavioral video acquisition. The oral cavity was imaged using a Blackfly S camera (BFS-U3-13Y3M-C, #36-451, FLIR, Edmund Optics) at 107 fps, corresponding to a temporal resolution of 9.35 ms per frame. The TTL signal and video-frame timestamps were acquired using a common temporal. All behavioral sessions were continuously recorded.

Jaw movement captured was quantified using DeepLabCut (DeepLabCut GUI 2.3.5)-based tracking of the upper and lower incisor tip to measure the amplitude of the mandibular deflection. The Euclidean distance between incisors was calculated and aligned to TTL-defined light onset; jaw-movement latency was resolved at the 9.35 ms frame interval.

### Anatomy

#### Brain tissue

Mice were deeply anesthetized and transcardially perfused with 30mL of ice-cold phosphate-buffered saline (PBS; pH 7.4), followed by 30mL of 4% paraformaldehyde (PFA) in PBS. Brains were removed with the brainstem intact and post-fixed overnight at 4 °C in 4% PFA. Tissues were then cryoprotected in 30% (w/v) sucrose in PBS at 4 °C until they sank, embedded in optimal cutting temperature compound (OCT; 4583, Sakura) and stored at −80 °C.

Brains were sectioned coronally at 50 μm using a Leica CM1950 cryostat. Serial sections were mounted on gelatin-coated slides (SouthernBiotech, SLD01-BX) and stored at −80 °C until further processing.

#### Trigeminal ganglia

Trigeminal ganglia were dissected after transcardial perfusion and post-fixed overnight at 4 °C in 4% PFA in PBS. Ganglia were cryoprotected in 30% sucrose in PBS at 4 °C until they sank, embedded in OCT, frozen on dry ice and stored at −80 °C. Tissues were sectioned at 20 μm using a Leica CM1950 cryostat. Serial sections were mounted on gelatin-coated slides and stored at −80 °C until staining.

#### Decalcification and sectioning of jaw tissue

Mandibles and maxillae were dissected after transcardial perfusion and post-fixed overnight at 4 °C in 4% PFA in PBS. Tissues were washed in PBS 3 times. Samples were decalcified for 7 d in 0.1 M EDTA (pH [value]) at 42 °C with agitation at 350 r.p.m. The EDTA solution was replaced every 2 d. After decalcification, tissues were cryoprotected in 30% sucrose in PBS at 4 °C for 3–5 d or until they sank.

Samples were embedded in OCT, frozen on dry ice and stored at −80 °C. Hemisected jaw preparations were sectioned at 50 μm using a Leica CM1950 cryostat. Serial sections were mounted on gelatin-coated slides and stored at −80 °C until staining.

### Immunohistochemistry and confocal imaging

Slide mounted tissue sections were equilibrated to room temperature and post-fixed in 4% paraformaldehyde (PFA) in phosphate-buffered saline (PBS) for 10 min. Sections were washed five times for 5minutes in PBS containing 0.1% Triton X-100 (PBST) and blocked for 30 minutes to 1 h at room temperature in PBST containing 10% normal donkey serum. Sections were incubated overnight at 4 °C with primary antibodies diluted in PBST containing 5% normal serum and antibody dilution buffer.

The following day, sections were washed five times for 5minutes in PBST and incubated for 2 h at room temperature with the appropriate species-specific secondary antibodies diluted in PBST containing 5% normal serum and antibody dilution buffer. Sections were protected from light during and after secondary-antibody incubation. After five additional washes in PBST, sections were coverslipped using ProLong Diamond Antifade Mountant (Thermo Fisher Scientific, P36970). Slides were allowed to cure for one day before imaging.

Images were acquired using an FV3000 laser-scanning confocal microscope (Evident Scientific) equipped with [4X, 10X, 40X objective lenses]. Single optical sections, serial z-stacks and tiled images were collected as appropriate for each analysis. For quantitative comparisons, laser power, detector gain, offset, pinhole size, pixel dimensions, z-step interval and objective settings were held constant across experimental and control samples within each experiment. Image acquisition and analysis were performed blinded to genotype or experimental condition. Images were processed using Fiji/ImageJ2 2.14.0/1.54j and/or Imaris 10.2 and any adjustments to brightness or contrast were applied uniformly across comparison groups.

### Antibodies

The following primary antibodies included: Chicken anti-GFP (1:500, yl4058591, Invitrogen), Sheep anti-GFP (1:500; # 163365, Bio-rad), Rabbit anti-DsRed (1:1000; 600-401-379, Rockland), Goat anti-Chat (1:500, AB144P, Millipore Sigma), Rabbit anti GFP-488 (A21311, Invitrogen).

The following secondary antibodies included:

Alexa Fluor 555 Goat anti-Rabbit (1:1000; #A32732, Thermo Fisher Scientific), and Alexa Fluor 488 Goat anti-Rabbit (1:1000; #A32731, Thermo Fisher Scientific). Alexa Fluor 647 Donkey anti-Sheep (1:500, 713-605-147, Jackson ImmunoResearch), Alexa Fluor 488 Goat anti Chicken (1:1000, ab150169, abcam), Alexa Fluor 647 Donkey anti-Goat (1:1000, a32849tr, Invitrogen)

### Specific controls

Antibody specificity was assessed using negative-control sections processed without primary antibody. These controls were incubated with secondary antibodies under otherwise identical conditions and imaged using settings matched to the corresponding experimental sections. Where applicable, reporter-negative tissue was also used to assess nonspecific immunoreactivity. No-primary-antibody controls showed no detectable signal under the acquisition settings used for experimental samples.

### Fluorescence *in situ* hybridization (FISH)

Fluorescence *in situ* hybridization was performed using third-generation hybridization chain reaction (HCR^TM^ v3.0 RNA-FISH; Molecular Instruments) according to the manufacturer’s protocol, with modifications from previously published methods (Ronan et al., 2025). All solutions and materials used before probe amplification were RNase-free. Fixed brainstem and trigeminal ganglion tissues were embedded in Tissue-Tek OCT compound (Sakura Finetek), cryosectioned at 50 μm for brainstem and 20 μm for trigeminal ganglia mounted gelatin-coated slides and stored at −80 °C until processing.

Sections were equilibrated to room temperature, post-fixed in 4% paraformaldehyde (PFA) in phosphate-buffered saline (PBS) for 15 min on ice and washed three times in PBS. Sections were then incubated for 10 min in PFA-chelating solution (triethanolamine; acetic anhydride). After pretreatment, sections were dehydrated through a graded ethanol series comprising 50%, 70% and 100% ethanol and were subsequently rehydrated.

Sections were equilibrated in HCR probe-hybridization buffer for 10 min at 37 °C and incubated with the appropriate HCR probe sets in probe-hybridization buffer for 24 h at 37 °C in a humidified chamber. Probe sets, initiator assignments, lot numbers and concentrations are provided upon request. After hybridization, sections were washed at 37 °C using the manufacturer’s probe-wash buffer and a graded series of probe-wash buffer and 2X saline–sodium citrate containing 0.1% Tween-20 (2X SSCT): 100%, 75%, 50% and 25% probe-wash buffer, followed by 2X SSCT. Sections were then washed in 2X SSCT at room temperature.

For signal amplification, fluorophore-conjugated HCR hairpins were prepared separately by heating each hairpin to 95 °C for 90 s and allowing it to cool to room temperature in the dark for 30 min. Sections were equilibrated in amplification buffer for 10 min at room temperature and incubated overnight at room temperature in amplification buffer containing the appropriate snap-cooled hairpins. Sections were subsequently washed in 2X SSCT and counterstained with DAPI (D9542, Millipore Sigma).

For imaging, sections were coverslipped using a mixture of 25% 2X SSCT with ProLong Diamond Antifade Mountant (Thermo Fisher Scientific, P36970). Images were acquired using a FV3000 laser-scanning confocal microscope (Evident Scientific) with acquisition settings held constant across samples included in the same quantitative comparison. For high-resolution imaging of presynaptic markers, images were acquired using a Nikon CSU W1-Sora confocal microscope (Nikon instruments). Acquisition settings were help constant across all samples within each quantitative comparison. Images were subsequently denoised and deconvolved using Nikon NIS-Elements software.

For sequential rounds of targeted multiplex RNA detection, sections were imaged after each amplification round and their positions were recorded. HCR probe and amplifier signals were removed by incubation in 75% formamide at 37 °C washed three times for 10 minutes. Loss of fluorescence from the preceding round was confirmed before subsequent hybridization.

Sections were then subjected to additional rounds of probe hybridization and HCR amplification using the procedure described above. Images from sequential rounds were registered using DAPI, native reporter fluorescence or anatomical landmarks. This approach was used for sequential detection of predefined transcripts rather than unbiased transcriptomic profiling.

#### Probe sets and cellular classification

HCR probe sets targeted *Fos, Calb2, Pdyn, Gad1, Slc17a6 (Vglut2), Slc32a1 (Vgat), Lhx1, Dlk1, Nts, SSt, Tac1, Slc17a7* was purchased from Molecular Instruments. Regions of interest were defined using Allen Mouse Brain Atlas/anatomical landmarks. A cell was classified as transcript-positive when prespecified criterion signal exceeding above background. Colocalization was scored only when transcript signals occurred within the same segmented cellular profile across the z-stack. Image acquisition and quantification were performed blinded to experimental condition.

### Slice electrophysiology

#### Acute brainstem slice preparation

Mice were deeply anesthetized with isoflurane and decapitated in accordance with the approved institutional protocol. The brain skull was rapidly removed and impersed in ice-cold modified artificial cerebrospinal fluid (aCSF) containing (in mM): 80 NaCl, 2.5 KCl, 1.25, NaH_2_PO_4_, 0.5 CaCl_2_, 3.5 MgCl_2_, 25 NaHCO_3_, 75 sucrose, 1.3 sodium ascorbate and 3.0 sodium pyruvate, with pH at 7.4. The solution was continuously equilibrated with 95% O_2_ and 5% of CO_2,_ with an osmolality of 310-320 mOsm.

Coronal brainstem slices (300 μm thick) containing (5Tr, 5Sol or the trigeminal motor nucleus) were prepared using a vibratome (Leica VT1200s). Slices were transferred and then incubated for about 40 minutes at 33 °C in oxygenated (95% O_2_ and 5% CO_2_) cutting solution which contains (in mM): 125 NaCl, 2.5 KCl, 2 CaCl_2_, 1 MgCl_2_, 1.25 NaH_2_PO_4_, 26 NaHCO_3_, 25 D-glucose, 1.3 sodium ascorbate and 3.0 sodium pyruvate, with pH at 7.2 and osmolality at 310-320 mOsm. After incubation, brain slices were placed in a recording chamber and perfused with oxygenated recording solution at a rate of 3 ml/minute at room temperature until recording.

#### Whole-cell patch-clamp recordings

For electrophysiological recordings, we performed whole-cell patch clamp recording. Neurons were visualized using an upright microscope. Patch pipettes were pulled from borosilicate glass capillaries (Sutter Instrument) and had resistance of 3-6 MΩ were filled with an internal solution that contains (in mM): 130 potassium gluconate, 5 KCl, 4 Na_2_ATP, 0.5 NaGTP, 20 HEPES, 0.5 EGTA, pH 7.28 with KOH adjustment, and measured osmolality at 310-320 mOsm.

Data were acquired by pClamp 10.0 software (Molecular Devices) with MultiClamp 700B patch clamp amplifier and digitized using a Digidata 1550B (Molecular Devices) interface. Responses were low pass filtered online at 2 kHz and digitized at 5 kHz. Series resistance, membrane resistance and capacitance were monitored throughout each recording. Light-evoked postsynaptic currents were recorded in voltage-clamp mode. Excitatory postsynaptic currents (EPSCs) were recorded at a holding potential of -70 mV, at which inhibitory currents were minimized, whereas inhibitory postsynaptic currents (IPSCs) were recorded at a holding potential of 0 mV, at which excitatory currents were minimized.

#### Optogenetic stimulation in acute brainstem slice

ChR2-expressing intradental afferent terminals were stimulated with 473-nm light, and ReaChR-expressing premotor axon terminals were stimulated with 595 nm light delivered through a pE-300 illumination system (CoolLED) and the microscope objective. Unless otherwise indicated, stimulation consisted of 1-ms pulses delivered at 5, 10 and 20 Hz. Optical output was calibrated to 5 mW before experiments 530-nm ReaChR experiments. Optical pulses were triggered through a Digidata 1550B digitizer using episodic protocols in pClamp 10.

For monosynaptic-connectivity assays, baseline light-evoked postsynaptic currents were first recorded in ACSF. Tetrodotoxin (TTX, 500 nM; Tocris Bioscience) was then applied to block action-potential-dependent transmission. After the light-evoked response was abolished, 4-aminopyridine (4-AP, 100 μM; MilliporeSigma) was co-applied in continued presence of TTX. Recovery of the light-evoked postsynaptic current in TTX plus 4-AP was taken as evidence supporting direct monosynaptic connectivity from the optogenetically stimulated presynaptic terminals.

#### Optogenetic circuit mapping of intradental afferent input to 5Tr and 5Sol neurons

To express ChR2 in tooth-innervating sensory afferents, AAV6.2/2-iCre (7.2 x 10E12 vg/ml, 21754, ETH VVF) was unilaterally injected into the right first molar (M1) of LSL-ChR2(H134R)-EYFP mice using the AAV-i.d. procedure described above. Acute brainstem slices containing the 5Tr or 5Sol were prepared 3 weeks after intradental viral delivery.

Whole-cell recordings were obtained from neurons located within the anatomically defined 5Tr or 5Sol. ChR2-expressing intradental afferent terminals were stimulated 473-nm light using 1-ms pulses at 5, 10 and 20 Hz. Light-evoked EPSCs were recorded at -70 mV under baseline conditions, after application of TTX, and during co-application of TTX and 4-AP. Abolition of the light-evoked EPSCs by TTX followed by recovery in TTX plus 4-AP supported direct monosynaptic excitatory input from intradental sensory afferents onto neurons in the 5Tr and 5Sol. Following electrophysiological recording, the cytoplasmic contents of individual responsive neurons were collected for single-cell RT-PCR to determine their molecular identities, as described below.

#### Optogenetic circuit mapping of Calb2+ 5Tr input to masseter motor neurons

To express ReaChR in Calb2+ 5Tr neurons, AAV9/2-dlox-ReaChR-mCitrine (5.9 x 10E12 vg/m, v261-9,VVF) was unilaterally injected into 5Tr of *Calb2*-Cre mice. Three weeks later, masseter motor neurons (5Ma) were retrogradely labeled by injection of the CTB-555 (cholera toxin subunit B conjugated to Alexa Fluor 555, (1%, 10.0mg/mL, 2.5uL, C34776, ThermoFisher) into the masseter muscles as described below. Acute brainstem slices containing the trigeminal motor nucleus were prepared 3-5 days after CTB injection.

Whole-cell recordins were obtained from CTB-555-labeled masseter motor neurons located ipsilateral or contralateral to the 5Tr viral injection. ReaChR-expressing Calb2+ axon terminals were stimulated during 530-nm light using 1-ms pulses. Light-evoked IPSCs were recorded at a holding potential of 0 mV under baseline conditions, after TTX application, and during co-application of TTX and 4-AP. Restoration of the light-evoked IPSCs in TTX plus 4-AP supported direct monosynaptic inhibitory input from Calb2+ 5Tr neurons onto masseter motor neurons. Ipsilateral and contralateral motor neurons were analyzed separately, and response in incidence was calculated as the number of light-responsive CTB-labeled neurons divided by the total number of recorded CTB-labeled neurons on each side.

#### Optogenetic circuit mapping of Pdyn+ 5Sol input to anterior digastric motor neurons

To express ReaChR in Pdyn+ 5Sol neurons, AAV9/2-dlox-ReaChR-mCitrine (5.9 x 10E12 vg/m, v261-9,VVF) was unilaterally injected into 5Sol of *Pdyn*-Cre mice. Three weeks later, CTB-555 was injected unilaterally into the anterior digastric muscle to retrogradely label its corresponding motor neurons. Acute brainstem slices containing the anterior digastric motor pool were prepared 3-5 d after CTB injection.

Whole-cell recordings were obtained from CTB-555-labeled anterior digastric motor neurons (5ADi). ReaChR-expressing Pdyn+ axon terminals were stimulated during 530-nm light using 1-ms pulses at 5, 10 and 20 Hz. Light-evoked EPSCs were recorded at a holding potential of -70 mV under baseline conditions, after TTX application, and during co-application of TTX and 4-AP. Restoration of LIGHT-EVOKED the EPSCs in TTX plus 4-AP supported direct monosynaptic excitatory input from Pdyn+ 5Sol neurons onto anterior digastric motor neurons (5ADi).

#### Retrograde labeling of jaw motor neurons

Mice were anesthetized with isoflurane. CTB conjugated to Alexa Fluor 555 (1%, 10.0mg/mL, 2.5uL, C34776, ThermoFisher) was injected into the masseter or anterior digastric muscle using a Hamilton syringe, volume and needle gauge. A total volume of 2.5 μl per muscle was delivered at the targeted muscle site. The needle was left in place for 5 min after injection before slow withdrawal. For anterior digastric labeling, CTB-555 was delivered to the anterior belly of the digastric while minimizing spread to surrounding tissues. Animals recovered for 3-5 days before acute brainstem slices were prepared.

#### Single-cell RT-PCR after electrophysiological recording

Following whole-cell recording, the cytoplasmic contents of individual neurons were aspirated into the patch pipette under visual control and collected for single-cell RT-PCR. PCR-amplified cDNA libraries were generated using the SuperScript IV Single Cell cDNA PreAmp kit (11752048, Thermo Fisher Scientific) according to the manufacturer’s instructions. cDNA qualify from each collected cell was assessed by PCR amplication of *Gapdh.* Recorded neurons were screened for *Calb2, Pdyn, Gad1, Vglut1, Lhx1, Dlk1, Nts* and *Gapdh* transcripts. Cells without detectable *Gapdh* amplification were excluded from molecular-identity analyses.

PCR reaction contained 1× reaction mix, 2 mM MgCl_2_, 250 μΜ each deoxynucleotide, 0.25 μM forward primer, 0.25 μM reverse primer, 2.5 U SuperScript One-Step RT-PCR (12594025, Invitrogen) and 1 μl template cDNA. PCR products were amplified for 35 cycles and separated on 1.5% agarose gels. Primer sequences are in Supplementary Table 1.

### Statistics and reproducibility

No statistical methods were used to predetermine sample sizes. Sample sizes were based on previous studies, preliminary experiments and conventions in the field. Statistical analyses were performed using GraphPad Prism 10 (GraphPad Software). All statistical tests were two-sided unless otherwise indicated. The statistical test used, definition of n, sample size and P values are indicated in the corresponding figure legends.

For comparisons between two independent groups, two-tailed unpaired Student’s t-tests were used. Paired two-tailed Student’s t-tests were used for paired measurements. Comparisons among more than two groups were performed using one-way ANOVA followed by the appropriate multiple-comparisons test, as indicated in the figure legends. For repeated measurements obtained from the same experimental unit, repeated-measures ANOVA was used where appropriate. Nonparametric tests were used when the assumptions for parametric testing were not met, as specified in the corresponding figure legends.

For electrophysiological experiments, the number of recorded cells and animals is indicated in the corresponding figure legends. Light-responsive cells were defined by the presence of reproducible light-evoked postsynaptic currents. For monosynaptic-connectivity experiments, synaptic-current amplitudes recorded under baseline ACSF, TTX and TTX plus 4-AP conditions were compared within the same recorded cells. Connection incidence was calculated as the number of light-responsive cells divided by the total number of recorded cells.

Data are presented as mean ± SEM unless otherwise stated. Statistical significance was defined as P < 0.05.

## Extended Data

**Extended Data Fig. 1.**
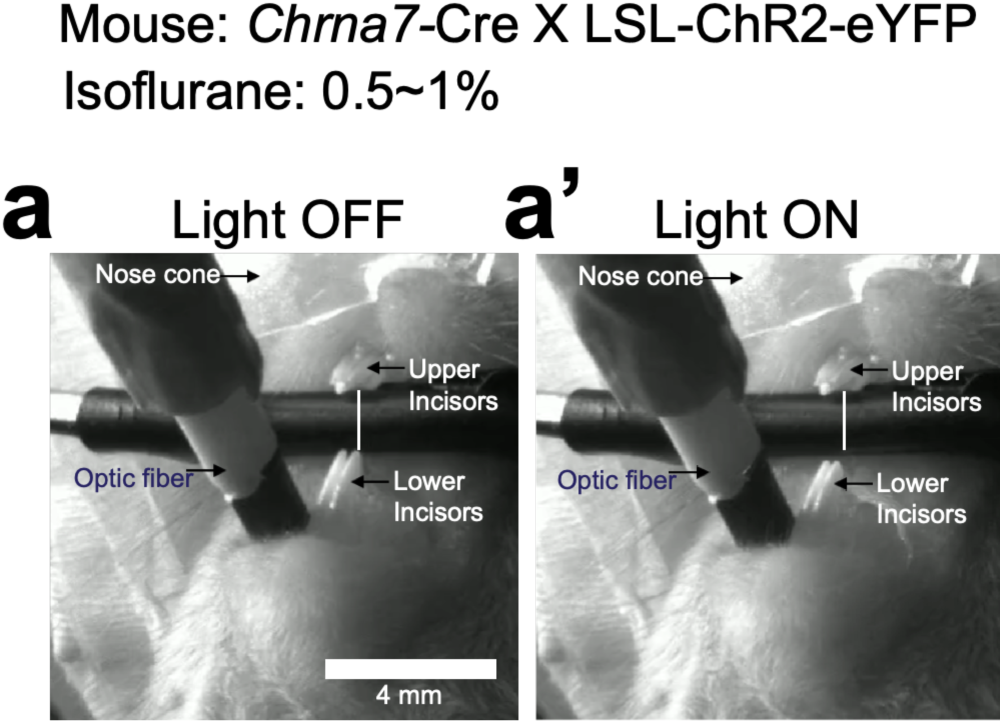
Molar stimulation in *Chrna7*-Cre; LSL-ChR2–eYFP mice evokes jaw opening. *Chrna7*-Cre mice were crossed with LSL-ChR2–eYFP reporter mice to drive ChR2–eYFP expression in *Chrna7*-expressing sensory neurons, including tooth-innervating sensory afferents. Blue-light pulses were delivered to the first maxillary molar (M1) over 1 min to activate dental sensory terminals and assess the resulting jaw-opening response. **a, a′,** Representative images acquired immediately before (**a,** light off) and during (**a′,** light on) molar photostimulation. In **a,** the white reference line connecting the upper- and lower-incisor tips indicates the baseline inter-incisor distance. In **a′,** an identical reference line was anchored to the upper-incisor tip at the same position. During photostimulation, the lower-incisor tip displaced away from the reference line, demonstrating an increase in inter-incisor distance consistent with jaw opening. Scale bar, 4mm. Representative responses were observed in stimulated *Chrna7*-Cre; LSL-ChR2–eYFP mice, n = 4 mice.

**Extended Data Fig. 2.**
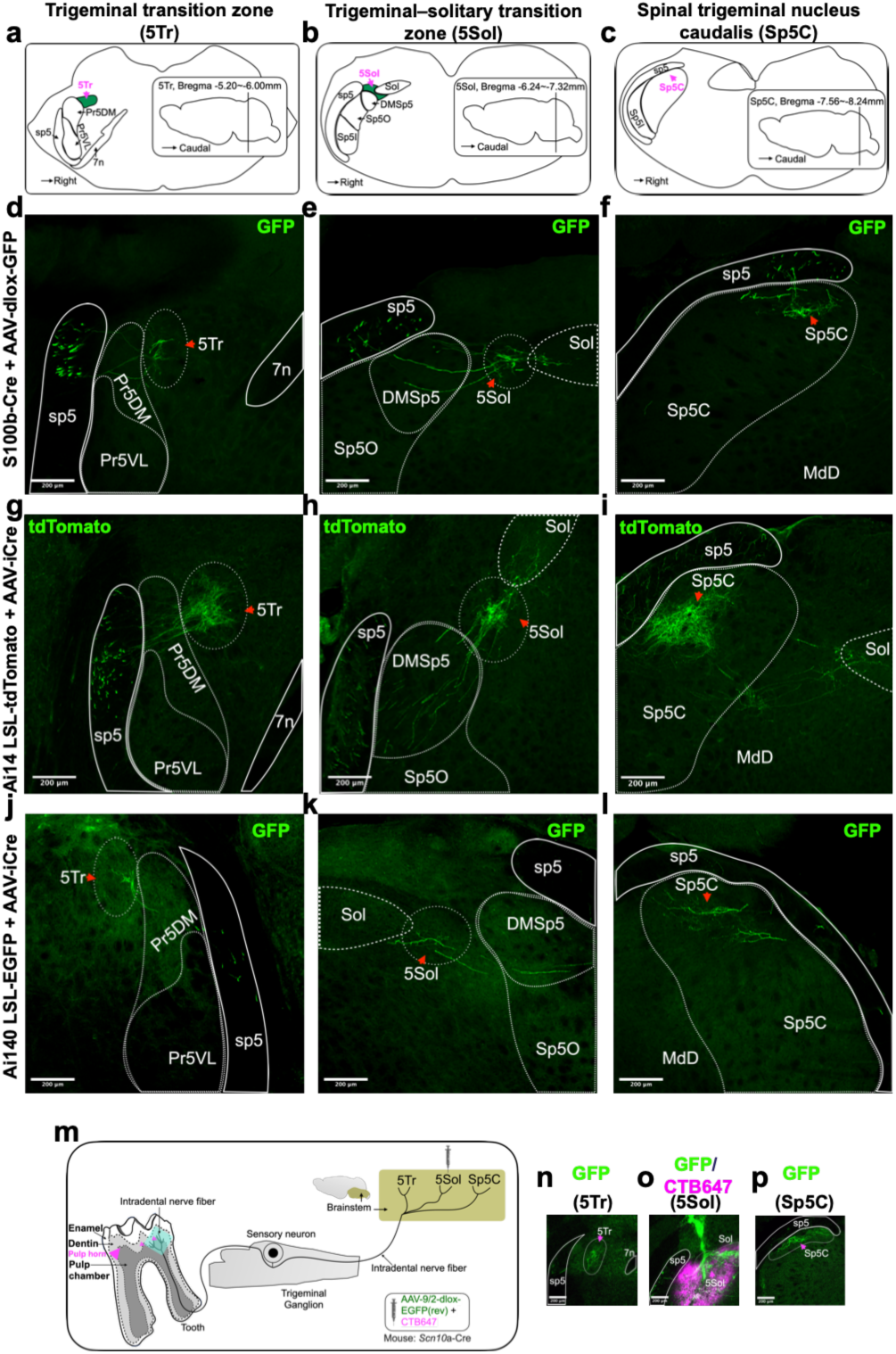
Complementary tracing strategies reveal distributed intradental afferent projections across three brainstem regions. **a-c,** Anatomical reference diagrams showing the rostrocaudal locations of the trigeminal transition zone (5Tr; **a**), trigeminal–solitary transition zone (5Sol; **b**) and spinal trigeminal nucleus caudalis (Sp5C; **c**). Regional boundaries were assigned with reference to the Allen Mouse Brain Atlas. **d-f,** Representative coronal brainstem sections from *S100b*-Cre mice following intradental delivery of AAV-dlox-EGFP. This strategy labeled tooth-innervating sensory neurons and their central axons. GFP-labeled afferent fibers were detected in 5Tr (**d**), 5Sol (**e**) and Sp5C (**f**). Arrowheads indicate representative projection fields. Scale bars, 200 μm. The labeling pattern was observed in n = 5 mice. **g-i,** Representative coronal brainstem sections from Ai14 LSL-tdTomato reporter mice following intradental delivery of AAV-iCre. Cre-mediated recombination labeled intradental sensory neurons and their central axons with tdTomato. TdTomato-labeled fibers, displayed in green pseudocolor, were detected in 5Tr (**g**), 5Sol (**h**) and Sp5C (**i**). Arrowheads indicate representative projection fields. Scale bars, 200 μm. The labeling pattern was observed in n = 3 mice. **j-l,** Representative coronal brainstem sections from Ai140 LSL-EGFP reporter mice following intradental delivery of AAV-iCre. EGFP-labeled intradental afferent fibers were detected in 5Tr (**j**), 5Sol (**k**) and Sp5C (**l**). Arrowheads indicate representative projection fields. Scale bars, 200 μm. The labeling pattern was observed in n = 5 mice. **m,** Schematic of the brainstem-targeted tracing strategy used to test whether 5Sol-associated intradental afferents also distribute projections to 5Tr and Sp5C. A viral tracer and CTB-647 were stereotactically co-injected into 5Sol, and CTB-647 was used to mark the injection site. **n-p,** Representative coronal brainstem sections from *Scn10a*-Cre mice following stereotactic injection of AAV-dlox-EGFP into 5Sol. GFP-labeled sensory fibers were detected in 5Tr (**n**), 5Sol (**o**) and Sp5C (**p**). CTB-647 signal (magenta) marks the 5Sol injection site (**o**). Scale bars, 200 μm. The labeling pattern was observed in n = 7 mice.

**Extended Data Fig. 3.**
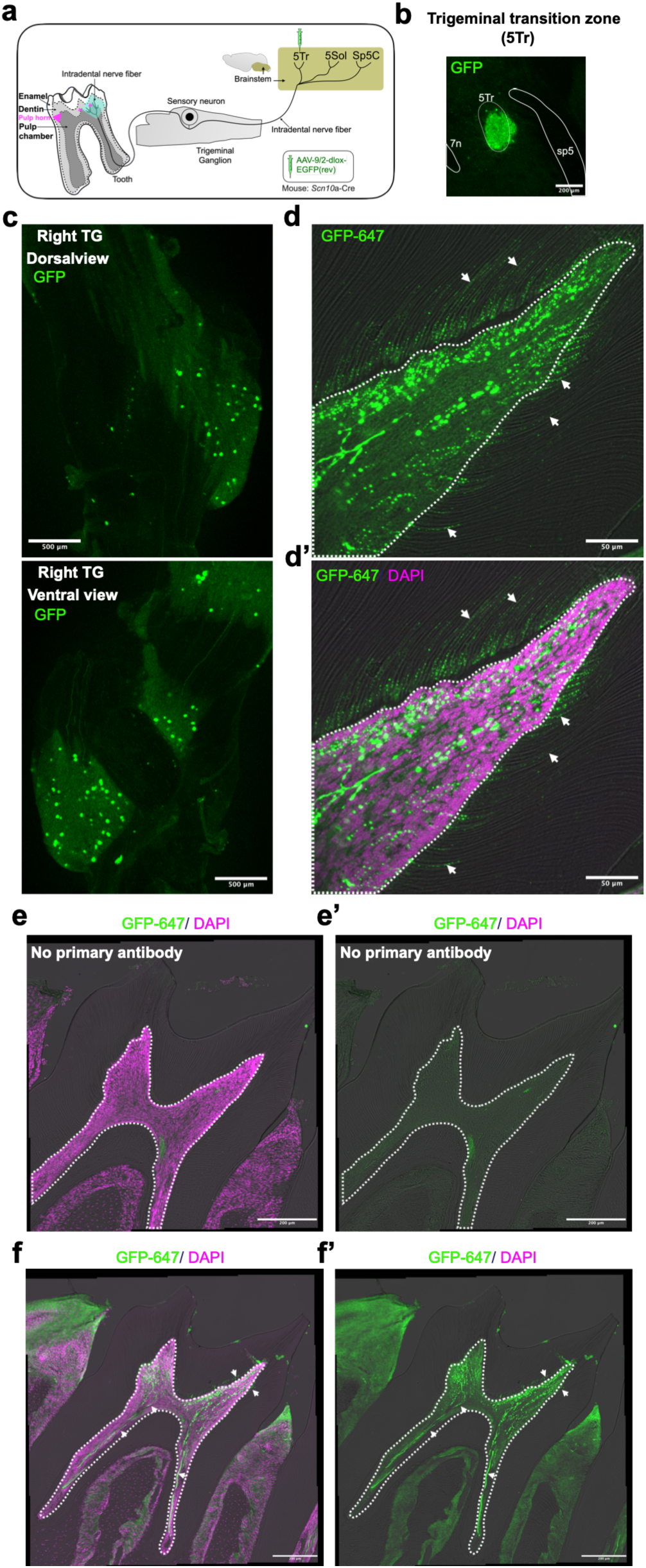
5Tr-targeted brainstem labeling identifies molar-innervating intradental sensory afferents. **a,** Schematic of the brainstem-targeted tracing strategy used to determine whether sensory afferents associated with the 5Tr projection domain innervate the molars. AAV-dlox-EGFP was stereotactically injected into the 5Tr of *Scn10a*-Cre mice, enabling Cre-dependent labeling of Scn10a-expressing neurons and their processes. **b,** Representative coronal brainstem section showing the 5Tr injection site and GFP-labeled sensory fibers following stereotactic delivery of AAV-dlox-EGFP. GFP-labeled fibers were also detected in 5Tr and Sp5C consistent with the distributed projection pattern identified by intradental tracing. **c,** Representative dorsal and ventral views of the trigeminal ganglion showing GFP-labeled sensory neuron cell bodies after 5Tr-targeted AAV delivery. GFP is shown in green. The labeling pattern was observed in n = 3 mice. **d, d′,** Representative high-resolution images of decalcified molar sections showing GFP-labeled sensory fibers within the dental pulp and extending toward the dentin (white arrows). GFP signal is displayed in green color, and DAPI-labeled nuclei are shown in magenta pseudocolor. White dashed lines delineate the pulp-dentin boundary. Labeled fibers were detected in the mandibular first molar (M1; **d, d′**) **e-f′,** Representative images of decalcified molar sections following 5Tr-targeted labeling. GFP-labeled sensory fibers were detected within the dental pulp and extended toward the dentin (white arrows, **f, f’**). White dashed lines delineate the pulp-dentin boundary. No GFP immunoreactivity was detected in negative-control sections processed without primary antibody (**e, e′**). Unilateral molar innervation was reproducibly observed in n = 3 mice. Scale bars, 200 μm (**b**), 500 μm (**c**), 50 μm (**d**), 200 μm (**e-f′**).

**Extended Data Fig. 4.**
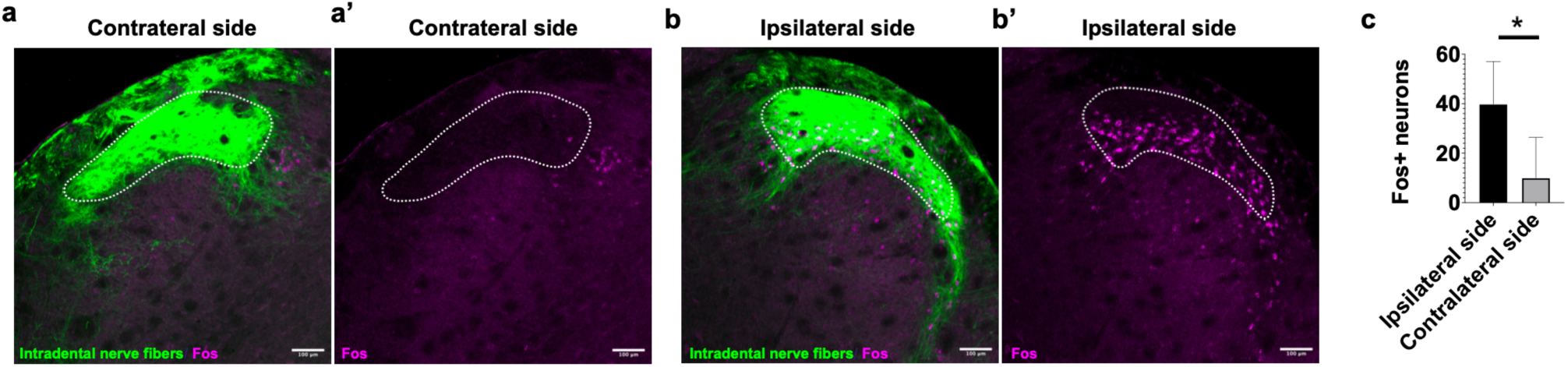
Optogenetic stimulation of the right molar preferentially induces *Fos* expression in the ipsilateral brainstem. **a-b’,** Representative coronal brainstem sections showing Fos induction after optogenetic stimulation of the right molar. Intradental sensory afferent fibers are shown in green and the white outlines delineate the brainstem Sp5C region used for quantification as region of interest (ROI). Representative images show the ipsilateral (**a, a’**) and contralateral side (**b, b’**) sides relative to the stimulated molar. Scale bars, 100 μm. Representative labeling patterns were observed in n = 5 mice. **c,** Quantification of Fos+ neurons in matched ipsilateral and contralateral brainstem regions. Right molar photostimulation produced greater Fos induction ipsilateral to the stimulated tooth (39.8 ± 7.7 Fos+ cells/ per ROI labeled by the white outline; 50 μm thickness) than on the contralateral side (10 ± 7.3 Fos+ cells/per ROI labeled by the white outline; 50 μm thickness). Data are presented as [mean ± SEM]; n= 5 mice. Statistical comparisons were performed using paired t-test, p-value = 0.023.

**Extended Data Fig. 5.**
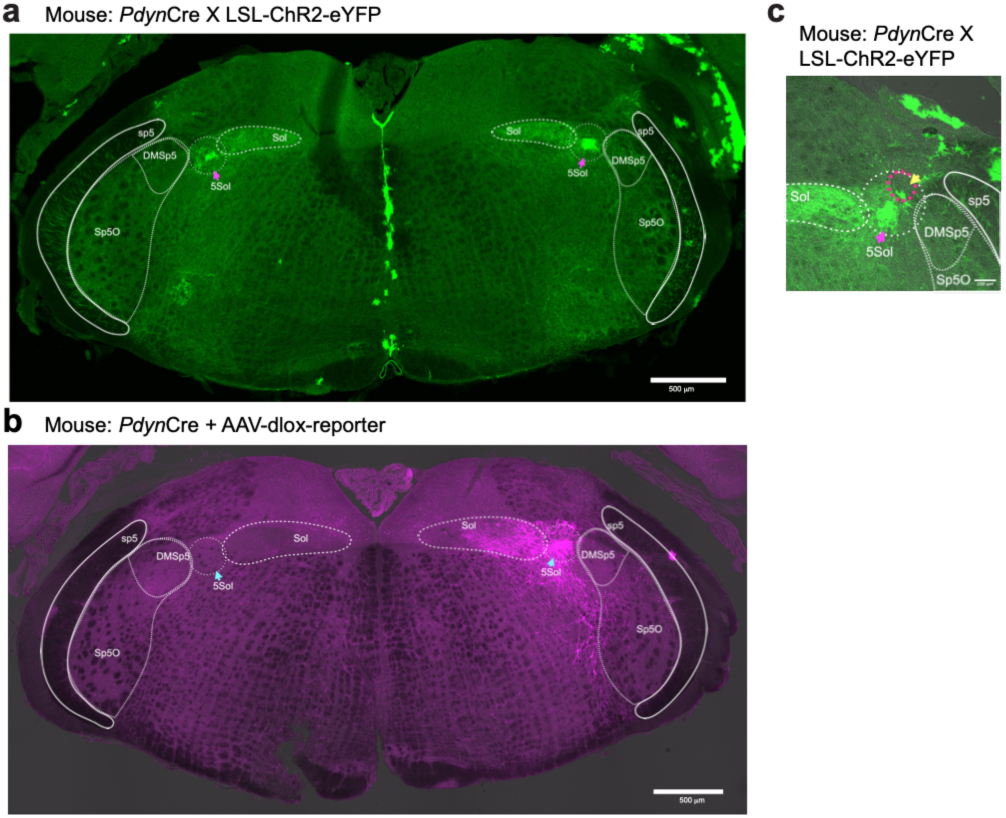
Verification of ChR2 expression and optic fiber placement in Pdyn+ 5Sol neurons. **a,** Representative coronal brainstem section from a *Pdyn*-Cre; LSL-ChR2-eYFP mouse showing Cre-dependent ChR2-eYFP expression in Pdyn-expressing neurons. Arrows indicate the bilateral 5 Sol regions. Scale bar, 500 μm. The expression pattern was confirmed in n = 6 mice. **b,** Representative coronal brainstem section from a *Pdyn*-Cre mouse following unilateral stereotactic injection of AAV-dlox-ChR2-mCherry into the right 5Sol. The injection was performed at the following coordinates relative to bregma: AP, -6.95 mm; ML, 1.30 mm; DV, -4.50 mm. ChR2-mCherry expression (magenta) was localized predominantly to the injected 5Sol region (arrow). Scale bar, 500 μm. Injection-site targeting was verified in n = 5 mice. **c,** Representative coronal brainstem section showing the optic fiber tip position relative to ChR2 expressing Pdyn+ neurons in the right 5Sol. The yellow arrow indicates the fiber tip location, and the magenta arrow marks the 5Sol region. Scale bar, 100 μm. Optic fiber placement was histologically verified in n = 5 mice

